# Interfacial vortex recapture enhances thrust in tiny water skaters

**DOI:** 10.1101/2024.06.17.599397

**Authors:** Pankaj Rohilla, Johnathan N. O’Neil, Paras Singh, Victor M. Ortega-Jimenez, Daehyun Choi, Chandan Bose, Saad Bhamla

## Abstract

Vortex recapture underpins the exceptional mobility of nature’s finest fliers and swimmers. Utilized by agile fruit flies and efficient jellyfish, this phenomenon is well-documented in bulk fluids. Despite extensive studies on organismal locomotion at the water’s surface, a vital fluidic interface where diverse life forms interact, hydrodynamics of interfacial vortex recapture remains unexplored. We investigate interfacial (on water) vortical hydrodynamics in *Microvelia americana*, one of the smallest and fastest water striders, skating at 50 body lengths per second (BL/s) or 15 cm/s. Their middle legs shed counter-rotating vortices, re-energized by their hind legs, demonstrating interfacial vortex recapture. High-speed imaging, particle imaging velocimetry, physical models, and CFD simulations show re-energization increases thrust by creating positive pressure at the hind tarsi, acting as a virtual wall. This vortex capture is facilitated by the tripod gait, leg morphology, and precise spatio-temporal placement of the hind tarsi during the power stroke. Our study extends vortex recapture principles from bulk fluids to the interface, offering insights into efficient interfacial locomotion, where surface tension and capillary waves challenge movement. Understanding interfacial vortex hydrodynamics can guide the development of energy-efficient microrobots to explore the planet’s water surface niches, critical frontlines of climate change and pollution.

**Significance Statement:** Interfacial Vortex Recapture in *Microvelia americana* extends the vortex recapture principles to the air-water interface, revealing an efficient locomotory mode in a challenging ecological niche. By demonstrating thrust enhancement through precise vortex interactions, our study bridges biology and fluid dynamics. This discovery informs the design of energy-efficient amphibious microrobots, capable of navigating the water interface with a new tripod gait paradigm, diverging from the conventional drag-based rowing designs. These findings are foundational for exploring and monitoring the water surface, an ecological interface vital for addressing climate change and pollution impacts.

---

**T**he unseen ballet of vortical forces orchestrates nature’s most efficient swimmers and fliers (1–7). These interactions, fundamental to minimizing energy expenditure and maximizing thrust, allow organisms to utilize energy from their own or others’ wakes (1, 2, 8–10). Jellyfish boost thrust by capturing vortices during relaxation, creating high-pressure zones (5, 11). Fruit flies capture leading-edge vortices during the fling motion, minimizing the energy required to generate new vortices (12, 13). Fish exhibit such efficient wake capture that even dead fish can swim upstream by resonating with oncoming Kármán street vortices (14, 15).

While these examples occur in bulk fluids, the air-water interface, a vital ecological niche, teems with life. From zooplankton, insects, and spiders to birds, reptiles, and plants, countless organisms interact at this boundary in marine and freshwater ecosystems (16–27). Despite the challenges of balancing surface tension, drag, buoyancy, and capillary waves, no documented examples of vortex recapture at this interface exist. Driven by curiosity about interfacial vortical interactions, we reveal a vortex re-energization mechanism in *Microvelia americana* (Hemiptera, Veliidae).

These millimeter-sized water walkers live on the water surface and are amongst the smallest and fastest in this ecological niche (body speed: **u**_*B*_ ∼ 50 BL/s, figure 2.d). Part of the infraorder Gerromorpha, they are found in creeks and ponds worldwide and include over 200 species (Figure S1) (28– 31). Unlike most water striders that use elongated middle legs for rowing, *Microvelia* employ all six legs to walk and run using a tripod gait (Figure 1.d, 2.a) (16, 17, 32, 33). Their unique morphology and kinematics enable them to recapture vortices shed from their middle legs, allowing them to speedily skate across the water surface. These amphibious insects, whose ancestors were terrestrial and used a tripod gait for movement on land, evolved to move on water while retaining this gait (16, 34–37). Using high-speed imaging, particle imaging velocimetry, physical models, and CFD simulations, we describe the interfacial vortex interactions during the water skating behavior of *Microvelia*.

**Fig. 1.**
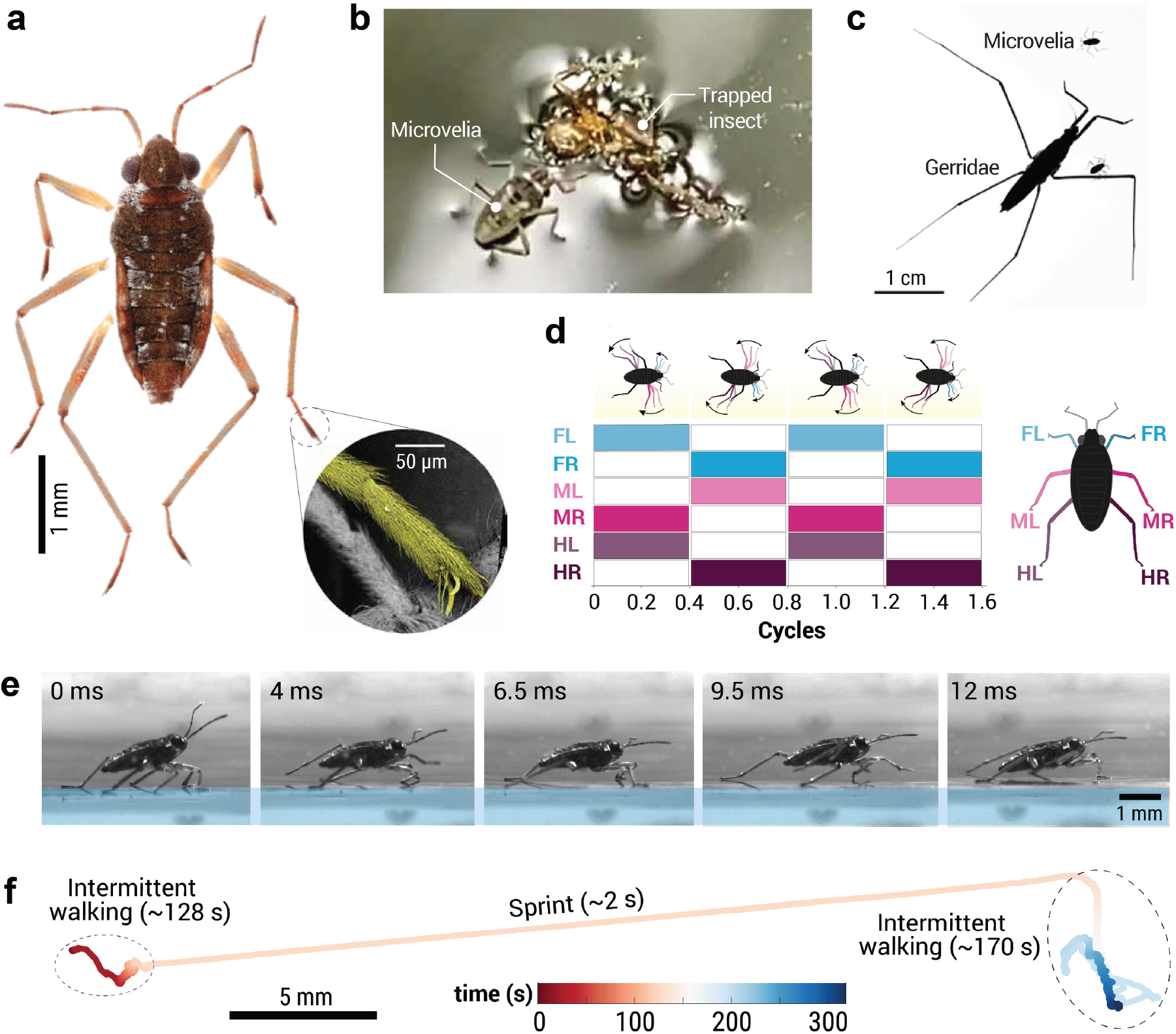
Behaviour and morphology of *Microvelia americana*. **(a)** Dorsal view of *Microvelia americana* with inset showing a SEM image of the dense hair coverage on middle leg tarsus (*pseudo-colored*). **(b)** *Microvelia Sp*. feeding on a trapped insect in a creek (Brunei), with legs deforming the water surface, forming dimples. **(c)** Size comparison showing *M. americana*’s small body size relative to commonly found water striders, *Gerridae*. **(d)** Alternating tripod gait plot for *M. americana* locomoting on the water surface, showing the gait cycle of each leg performing power (color filled boxes) and recovery strokes (empty boxes). **(e)** Snapshots showing the side view of *M. americana* walking on water. **(f)** Dynamics of *M. americana* on water, indicating short skating escape-sprints (∼2 s) and intermittent walking behavior over a 5-minute period.

**Fig. 2.**
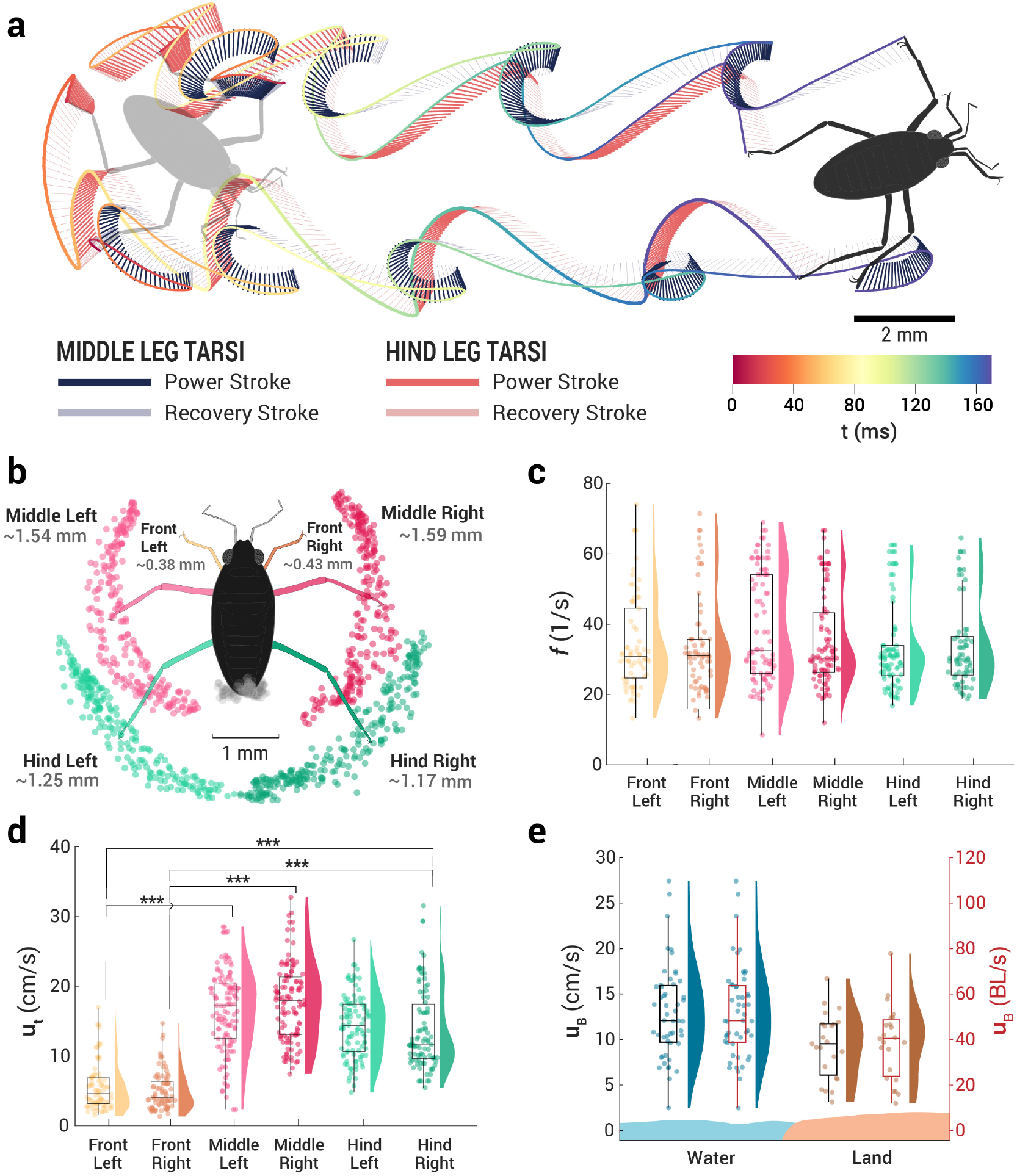
Interfacial kinematics of *Microvelia*. **(a)** Tarsal trajectories of middle and hind legs of *Microvelia*. The solid lines represent the power strokes, while the faded blue and red lines show the recovery strokes. The trajectories illustrate the time spent by the tarsi during movement. **(b)** Stroke amplitudes of the middle and hind legs (n = 15), illustrated with their tarsal tip trajectories relative to the motion of their respective shoulder joints. The middle legs exhibit larger stroke amplitudes (*λ*_*ML*_ ∼ 1.54 ± 0.43 mm and *λ*_*MR*_ ∼ 1.59 ± 0.74 mm) compared to the hind legs (*λ*_*HL*_ ∼ 1.25 ± 0.46 mm and *λ*_*HR*_ ∼ 1.17 ± 0.47 mm). **(c)** Stroke frequency (N = 3, n = 15) of the middle and hind tarsi, showing an average stroke frequency of *f* ∼ 30 strokes/s. **(d)** Peak tarsi speeds (**u**_**t**_) of *Microvelia* during power strokes on water. The middle legs achieve higher peak linear speeds during power strokes (∼17 cm/s) compared to the hind legs (∼ 14 cm/s). This indicates that the middle legs act as the main hydrodynamic thrust propulsors, with higher acceleration (∼ 2500 cm/s^2^) compared to the hind legs (∼ 2000 cm/s^2^). **(e)** Body speed (**u**_**B**_) of *Microvelia* on water and land (styrofoam) in cm/s (left Y axis) and BL/s (body lengths per second, right Y axis). The average maximum body speed on water is ∼ 15 cm/s (∼ 50 BL/s), compared to ∼ 10 cm/s (∼ 40 BL/s) on land. For (**c-e**), semi-violin plots show the distribution of the kinematics data, overlaid with jitter data points, while the box-and-whisker plots indicate the median and quartiles (25%, 50%, 75%, and 100%).

## RESULTS

### Skating on water

*Microvelia* possess dense hair coverage on their bodies and legs (Figure 1.a) (38). SEM analysis reveals a tarsal hair density of ∼ 15, 000 hairs/mm^2^ (*n* = 3), comparable to *Velia caprai* and *Gerridae*. (16, 39, 40). This dense coverage enables *Microvelia* to maintain a Cassie-Baxter state (41), limiting water infiltration and maintaining superhydrophobicity, leading to dimples at air-water surface contact points (Figure 1.b). The low Weber number, *We* = *ρv*^2^*l/σ* ≪ 1 (see Table S1) indicates that surface tension forces dominate over inertial forces in their interfacial locomotion, similar to other water striders like *Gerridae* (24).

Unlike water striders such as *Gerridae* that use a rowing gait, *Microvelia* employ an alternate tripod gait typical of terrestrial insects. In this gait, at least three legs – the front leg (FL), the contralateral middle leg (ML), and the ipsilateral hind leg (HL) – perform a power stroke on water (Figure 1.d,e), while the other legs recover in air or sometimes on water (SI Video 1).

To understand their interfacial locomotion behavior, we examine their dynamics over a 5-minute period in the lab. During this time *Microvelia* primarily engage in intermittent walking, spending 99.6% of the time in this mode. However, they occasionally sprint as an escape response, skating a distance of ∼ 30 mm in ∼ 2 seconds (Figure 1.e). The temporal trajectory of the middle and hind legs shows overlapping paths during this skating mode, leading us to the hypothesis of interfacial vortical interactions (Figure 2.a).

During the skating mode, the middle legs of *Microvelia* act as the main hydrodynamic thrust propulsors (32, 33). These legs exhibit a stroke amplitude 23% larger than the hind legs, while maintaining the same stroke frequency (Figure 2.b,c). This larger stroke amplitude allows for greater body displacement with each stroke, enhancing thrust. The middle legs achieve higher average peak linear speeds during power strokes, ∼21% faster than the hind legs (Figure 2(d)). This increased speed, combined with about 25% greater acceleration, indicates their dominant role as primary thrust generators (33). The tarsal speeds of the front legs are statistically lower than those of both the middle (*p <* 0.001) and hind legs (*p <* 0.001). However, due to high variability, the difference between the tarsal speeds of middle and hind legs is not statistically significant (*p >* 0.5). Additionally, Microvelia can traverse on both water and land with comparable body speeds with no statistical difference (Figure 2(e), *p >* 0.5).

### Interfacial hydrodynamic interactions

During the power stroke, the middle leg tarsi shed pairs of counter-rotating vortices (Figure 3.a, stage I). These vortices travel downstream, interacting with the hind tarsi, which enter the water at various spatio-temporal locations. The front tarsi generate weak vortices that dissipate without interacting with other tarsi (SI Video 2).

**Fig. 3.**
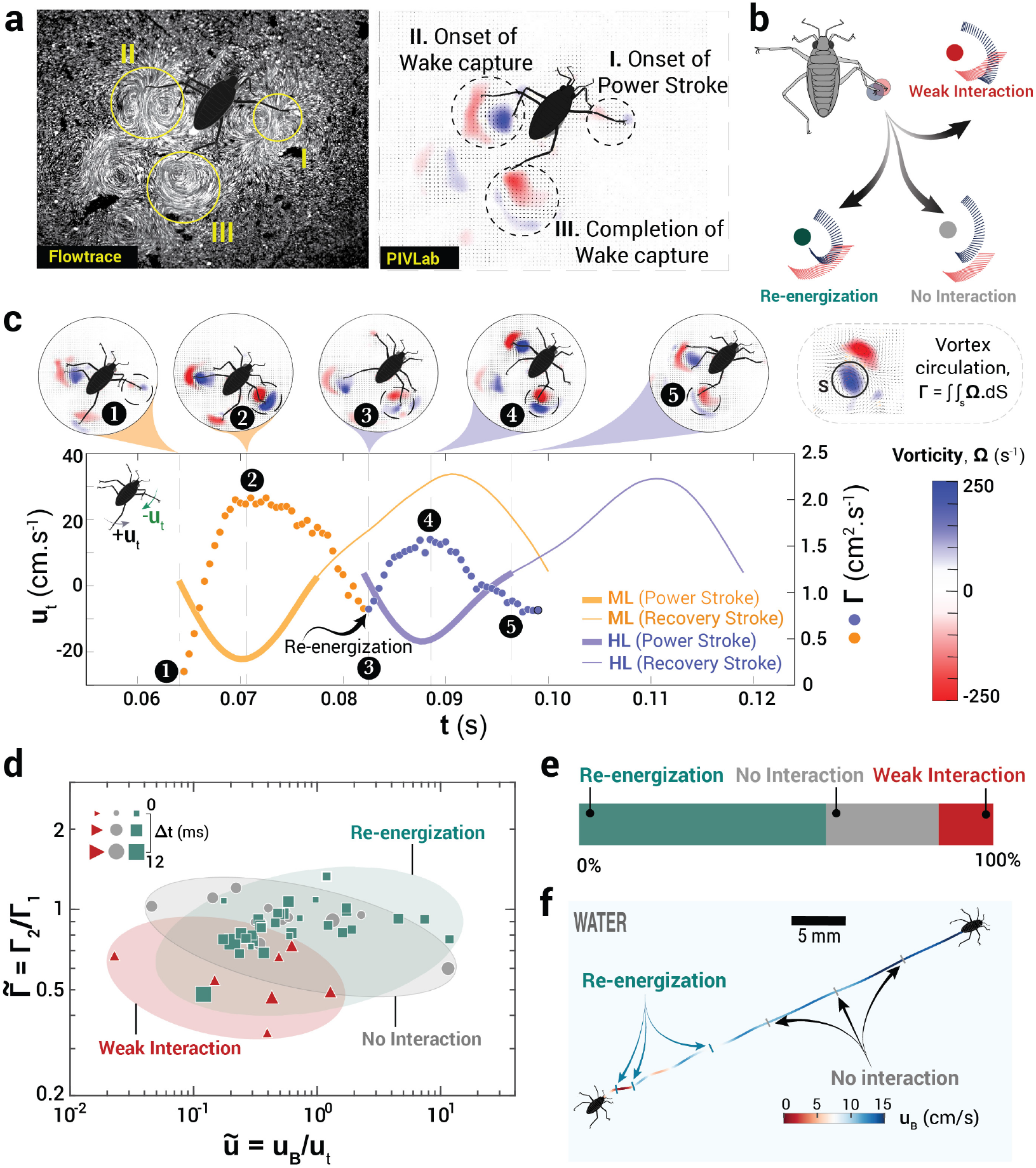
Hydrodynamic interactions in *Microvelia*’s interfacial locomotion. **(a)** Stages of vortical shedding from the power strokes of the middle tarsus and their subsequent interactions with the hind tarsus. LHS: Flowfield streamlines visualization in Flowtrace (42) and RHS: vorticity field generated in PIVlab (43). Stage I - Vortices generated during the onset of the power stroke of the middle right tarsus, (II) Hind legs stepping into the vortices shed from the middle left tarsus, (III) re-energized vortices from the hind right tarsus; LHS shows the vorticity field corresponding to the frame on the right. **(b)** Illustrations represent the three different outcomes of vortical interactions based on the trajectory of the hind and middle tarsi. **(c)** Representative tarsal velocity profiles of the middle-right and hind-right tarsi of *Microvelia* walking on water and the corresponding circulation (filled circles) of the vortices for the case of vortex re-energization. **(d)** Effect of normalized body speed (relative to hind tarsi speed) on the circulation ratio of vortices originating from the middle legs pre- and post-interactions with the hind tarsi. **(e)** Percentage outcomes of the vortical interactions of the hind tarsi with vortices shed from the middle tarsi, and **(f)** Different vortical interactions within a single run on water in *Microvelia*.

The exact location and timing of the incident hind tarsi relative to the vortices dictate the outcome of these interactions. Favorable interactions for vortex re-energization occur when the hind tarsi enter parallel to the downstream of the middle tarsi’s vortex pair, positioned to sweep them (Figure 3.a, stage II and III). Body rocking and turning can misalign these interactions, altering the hind legs’ angle of attack and leading to weak vortical interaction or no interaction (Figure 3.b). Additionally, if *Microvelia* moves at high speed, its body can pass over the middle leg vortices before the hind legs can interact with them, emphasizing the importance of timing (Figure 3.f).

We measure the circulation of vortex pairs generated by the middle tarsi during re-energization until they dissipate after hind tarsi interaction. Circulation, Γ = ∫ ∫_s_ Ω.*dS*, where Ω is the vorticity and *S* is the bounded area, measures the vortices’ strength. As the middle leg initiates the power stroke (Figure 3.c, point 1), the vortices’ circulation increases, peaking at Γ = 2 cm^2^/s (t = 71 ms), corresponding to the maximum tarsal speed (22 cm/s, t = 70 ms). The middle leg then decelerates, reducing Γ as the vortices dissipate (point 3). The hind tarsi then enters the wake, re-energizing the vortices to enhance the circulation to a second, lower peak of Γ = 1.6 cm^2^/s (t = 88.5 ms) due to a lower hind-tarsal speed of 17 cm/s (Figure 3.c, point 4). This cycle ends with the hind tarsi completing their power stroke and dissipating the vortices (SI Video 2, Figure S3).

Across 52 instances of hind tarsi interacting with vortex dipoles from the middle tarsi (6 specimens), hind tarsi re-energized 60% of the vortices, while not interacting with 27% of the vortices, and the remaining cases exhibited weak interaction (Figure 3.e). We compare the normalized peak circulation (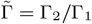, where Γ_2_ and Γ_1_ are the circulation of the middle leg vortices before and after their hind tarsal interactions) with normalized body speed (**ũ** = **u**_*B/*_**u**_*t*_) and the time interval (Δ*t*) between strokes (Figure 3.d). Vortical re-energization primarily occurs when the hind tarsi slaloms in between the slow moving vortices from middle tarsi, with shorter Δ*t* (typically *<* 6 ms), during the initial acceleration phase of the skating sprint (Figure 3.f). The slaloming of hind tarsi in between the bipolar vortices occurs due to the natural arc-like path of tarsi. At higher body speeds, longer Δ*t*, or due to body turning or rocking, the hind tarsi miss the vortices resulting in no interaction. When the hind legs skate across the pair of vortices rather than slaloming between them, the interactions tend to weaken the vortices, leading to weaker vortical interactions (Figure 3.d). Collectively, this reinforces that both the hind tarsi’s entry position relative to the middle tarsi (angle of attack) and the inter-stroke interval play critical roles in determining the outcome of these interactions. Although top-view planar interfacial PIV did not show the presence of capillary waves, shadowgraphy imaging enabled us to visualize these waves. The power strokes of the middle tarsi generate capillary waves, which dissipate before the hind tarsi begin their power stroke (SI Video II). Note that no noticeable capillary waves were observed during hind tarsi’s power stroke, suggesting that capillary waves have minimal to no impact on interfacial vortical interactions.

### Interfacial vortical recapture increases thrust in *Microvelia*

Reconstructed pressure fields from PIV-measured velocity fields reveal insights into vortical interactions with the hind tarsi of *Microvelia* (Figure 4.a). During vortex re-energization, a local pressure gradient forms from upstream to downstream of the hind tarsi, generating the highest relative pressure (Δ*p* ∼ 5 Pa). In contrast, weak vortical interactions results in lower relative pressure (Δ*p* ∼ 2 Pa, Figure S6), with cases of no interaction showing similarly low pressure.

**Fig. 4.**
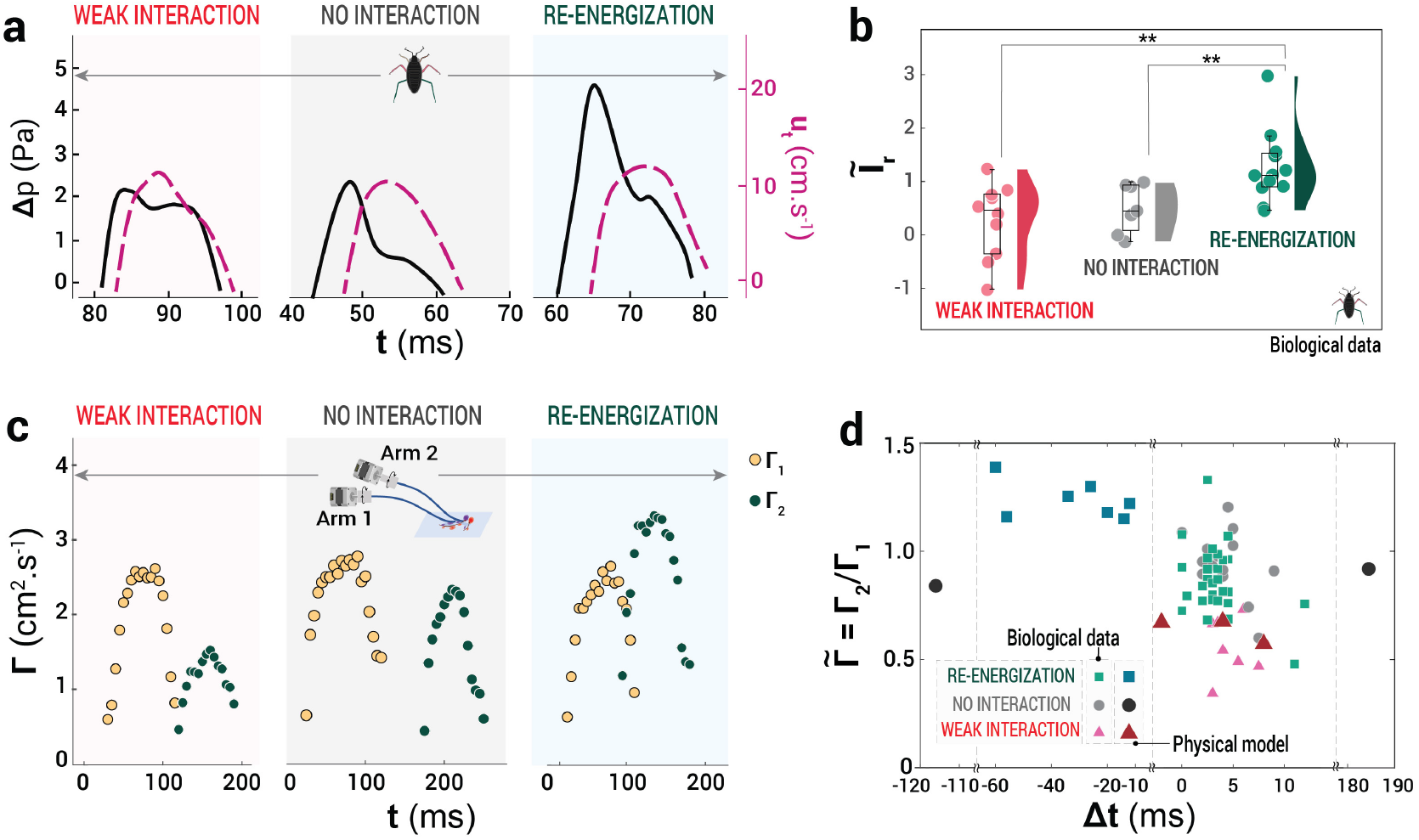
Quantifying interfacial vortical interactions in *Microvelia* and the physical model. **(a)** Temporal evolution of relative pressure (**ΔP**) and tarsal speed (**v**_*t*_) of the hind leg. No-interaction and weak interaction cases represent the hind left tarsi, while re-energization corresponds to the hind right tarsi. **(b)** Normalized impulse for different types of vortical interaction. The semi-violin plot shows the distribution of the data as a jitter plot, while the box and whisker plot represent the median and the four quartiles (25%, 50%, 75%, and 100%) for normalized impulse data. *Microvelia* specimens (N=8) and number of vortical interactions (*n*_*i*_ = 29). Normalized impulse for re-energization case was significantly higher than that of the no-interaction case (*p <* 0.01) and the weak interactions case (*p <* 0.01). Wilcoxon rank-sum test was used to perform the statistical analysis. **(c)** Temporal evolution of the vortex circulation **Γ** for each robotic arm with varying Δ*t* showing different vortical interaction outcomes. **(d)** Regime map of normalized circulation (**Γ**_**2**_ */***Γ**_**1**_) for varying Δ*t*. **Γ**_**1**_ and **Γ**_**2**_ represent peak circulation from the middle leg (or first arm) and hind leg (or second arm), respectively. For the biological data, the number of specimens is N = 7, with 53 vortical interactions, while the physical data show 12 vortical interactions.

We calculate the total impulse by integrating the relative pressure over time, **I** =∫_*T*_ Δ*pAdt*, where *T* is the duration of the power stroke and *A* is defined as the constant interfacial area between the water surface and the tarsi, approximated as constant value, *A* = *Ld*, where L is the tarsi length and d is the submergence depth. Normalizing the impulse 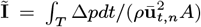, isolates the impact of hind tarsal interaction from tarsal speed. Excluding the impulse from the middle tarsi yields the relative impulse, **Ĩ**_*r*_ = **Ĩ***/* **Ĩ**_*middle*_. Here, the pressure analysis includes inherent numerical errors amplified by the second-order finite difference method. Additionally, three-dimensional measurements will be required for a more precise estimation of impulse generation.

Our results show that vortex re-energization produces a normalized impulse (**Ĩ**_*r*_ ∼ 1.26), ∼2.5 times higher than no interaction case (**Ĩ**_*r*_ ∼ 0.50) and ~ 4.6 times higher than the weak interactions case (**Ĩ**_*r*_ ∼ 0.27) (Figure 4.b). This increased impulse results from enhanced fluid entrainment during re-energization, which raises pressure in the tarsal plane. When hind tarsi step into the center of the vortex pair, they entrain more fluid mass due to the converging flow driven by the vortical motion (44), leading to increased pressure and greater thrust.

The observed rise in normalized impulse during re-energization illustrates *Microvelia*’s ability to harness energy from its own wake, a phenomenon we call ‘Interfacial Vortex Recapture’. Typically, wakes signify lost energy to the environment. By stepping into vortices generated by its middle legs during previous strokes, *Microvelia* harnesses this energy to increase thrust production by the hind legs. This mechanism, driven by its tripod gait and interfacial movement, enables *Microvelia* to generate thrust at the air-water interface effectively.

### Physical models validate inter-stroke interval in interfacial vortical interactions

To evaluate the effect of inter-stroke intervals (Δ*t*) on vortical interactions, we use a physical model. The model simulates *Microvelia*’s middle and hind tarsi power strokes on water, varying Δ*t* to alter the hind tarsi’s angle of attack to the vortices shed by the middle legs. The maximum speeds of the wires of arm 1 and arm 2 skating across the water surface were 11.4 cm/s and 8.14 cm/s, respectively. The Reynolds number of the physical model’s arms (*Re* ∼ 8 − 11) was within the range of *Microvelia*’s tarsal motion (*Re* ∼ 2 − 21, see Table S1). The first arm generates a counter-rotating vortex dipole, which the second arm interacts with, depending on Δ*t* (Figure 4.c).

For large Δ*t* = −116 ms, the first arm’s vortices dissipate before the second arm’s entry, resulting in no interaction (SI Video 3, Figure S7). Reducing the time interval ( −60 *<* Δ*t <* −10) allows for re-energization, with the second arm’s vortices showing higher normalized circulation 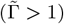 (Figure 4.d). However, at very short intervals ( −10 *<* Δ*t <* 10 ms), capillary waves generated by the arms disrupt the vortices, leading to lower circulation 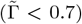. Further increase in Δ*t* beyond 10 ms resulted in collision of the robotic arms of the physical model (see Figure S7). This limitation of the physical model limited the collection of the data in this space Δ*t >* 10 ms.

Although the Reynolds numbers of the physical model’s arms were matched to the tarsal motion of *Microvelia*, the trajectories differ: the robotic arms followed a fixed path with same speed, whereas *Microvelia* exhibited stroke-to-stroke variability in tarsal trajectories and speed. This variation, combined with differences in Δ*t*, led to normalized circulation data that did not align with the regimes observed in the physical model for different types of vortical interactions (Figure 4.d). Despite these discrepancies, normalized circulation provides insight into both systems, demonstrating that optimal inter-stroke intervals can enhance vortex circulation 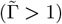, leading to effective vortex re-energization. In *Microvelia*, however, re-energization does not always correspond to 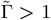, due to variations in tarsal speed, Δ*t*, and the trajectories of the middle and hind legs (Figure 2.e).

### Computational analysis of thrust enhancement during vortex re-energization: single-phase simulations

First, we perform two-dimensional (2D) bulk flow simulations using a canonical model of two rectangular plates that undergo prescribed rotation and translation to demonstrate thrust enhancement via vortex capture by the hind (second) plate. We model *Microvelia*’s tarsi as rectangular plates representing the top view of a cylinder, with an aspect ratio of 20, defined by the ratio of length *L* and width *L/*20. Simulations were performed within a low Reynolds number regime (*Re* = 20), consistent with the *Re* range observed in *Microvelia*’s motion (*Re* ∼ 2 − 21). We use the plate length and its translational velocity as the characteristic length and velocity scales, respectively.

We use rectangular geometries to represent tarsi in single-phase 2D simulations, aiming to capture the interfacial vortex capture mechanism seen from the top view in our planar PIV experiments. Mimicking the physical model kinematics, the first plate rotates counterclockwise, and the second rotates clockwise, initiating motion after a time gap (Δ*t*) to traverse the vortical wake of the first plate (see Figure 5(a), SI Video 4). The computational domain spans a height and width of 30*L*, and Figure S12(a) in the supplementary material shows a zoomed-in view of the mesh. These simulations evaluate the role of vortex re-energization on thrust based on the robotic arms’ trajectories rather than replicating the exact kinematics of *Microvelia*.

**Fig. 5.**
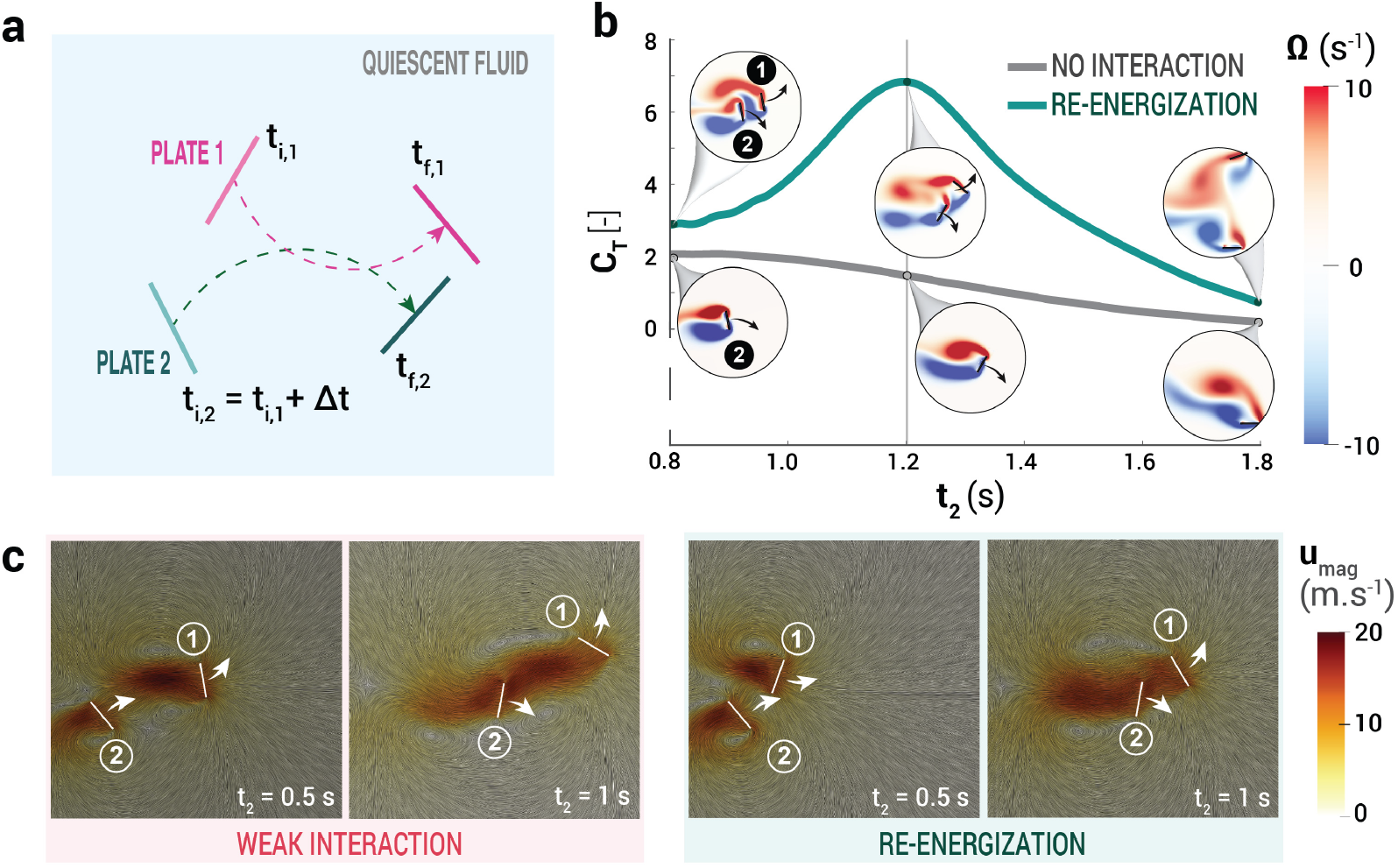
Single-phase 2D CFD analysis of vortical interactions. **a)** Illustration of the trajectories of the two plates used in 2D CFD simulations in quiescent water. **b)** CFD results show the effect of the time interval between plate movements (in quiescent fluid) on vortical interactions depicted by velocity magnitude contours. The second plate starts moving at *t* = 0 s with Δ*t* = 0.2 s for re-energization and Δ*t* = 0.5 s for weak interaction. In snapshots at *t* = 1 s, arrows indicate the enhanced and reduced velocity field due to wake capture and weak vortex interaction, respectively. **c)** Temporal evolution of the coefficient of thrust (*C*_*T*_) of the second plate for re-energization and no interaction. Snapshots show the interaction’s impact on instantaneous vorticity fields at different times.

The temporal evolution of thrust coefficient (*C_T_*) for the second plate (Figure 5(b)) shows both the vortical re-energization and no-interaction cases, with corresponding vorticity contours shown in the insets. We define *C_T_* as 2*T/*(*ρ*_*f*_**u**_**r**_^2^*A*), where *T* is the thrust force, *ρ*_*f*_ is the fluid density, **u_r_** is the relative velocity experienced by the second plate, and *A* is its projected area. The temporal variation in *C_T_* reflects the changing vortex–plate interactions over time. For Δ*t* = 0.2 s, *C_T_* peaks at *t* = 1.2 s, when vortex dipoles from both plates interact and amplify circulation. As the plates move apart, *C_T_* decreases, indicating reduced wake interaction. The observed thrust enhancement through wake capture corresponds with variations in fluid impulse governed by circulation and vortex core velocities. This unsteady flow behavior is consistent with pressure data from *Microvelia* and the physical model (Figure 4), where optimal stroke timing increased entrainment and thrust, demonstrating enhanced thrust through vortex re-energization.

Contours of flow velocity magnitude and streamlines, visualized using line integral convolution of the velocity field, show the flow generated by two plates moving through quiescent fluid with two different time intervals between the onset of their motions (Δ*t*); see Figure 5(c). The flow velocity magnitude behind the plate is as a key indicator of the thrust it generates. The vortical interactions change significantly with varying Δ*t*, consistent with observations from the physical model experiments. For Δ*t* = 0.2 s, the second plate captures the first plate’s wake, entering its recirculation region closely (SI Video 4). This leads to the merging of vortex cores with the same sense of rotation, increasing overall circulation and enhancing thrust. In contrast, for Δ*t* = 0.5 s, the second plate weakly interacts with the first plate’s wake from the first, resulting in lower flow velocities, and reduced thrust. The magnitude of flow velocity (**u**_*mag*_), particularly in the wake region, directly correlates with the momentum transfer from the plate to the fluid. The higher wake velocities for Δ*t* = 0.2 s indicate stronger momentum transfer and greater thrust, driven by a larger velocity difference between the fluid and the plate, which leads to a higher pressure differential.

The primary goal of these idealized simulations in single-phase flow is to isolate and understand the role of vortex interactions, from the top view as in planar PIV, in enhancing the thrust generated by the second plate as it traverses through the wake of the second plate. However, these simulations do not capture the full complexity of *Microvelia*’s tarsal kinematics, particularly the tarsal entry from air to water surface.

### Computational analysis of thrust enhancement during vortex re-energization: two-phase simulations

Next, to incorporate the effect of surface tension on the vortex capture mechanism at the air-water interface, we perform 2D two-phase flow simulations from a side-view perspective, where the circular cross-section of the tarsus grazes the interface. In this case, we used a Reynolds number of 50, based on the cylinder diameter (*d*), which is of the same order of magnitude (*Re* ∼ 𝒪 (10)) as the tarsal motion of *Microvelia*, the physical model, and the bulk-phase simulations. The computational domain for these simulations spans a length of 40*d* and a width of 20*d*, with a circular body representing the cross-section of the *Microvelia* tarsus. The mesh topology used in these simulations is presented in Figure S12(b).

We selected the grid size for the two-phase simulations through a grid sensitivity study using the Richardson extrapolation method (46), with the results shown in Figures 6(e)–6(f) and Table S2. We consider three different grid cell sizes: mesh 1 (126,600 cells), mesh 2 (253,200 cells), and mesh 3 (506,400 cells). For each mesh, we compute the mean value of *C_T_* and the peak Γ for the two-cylinder case at *Re* = 50 with a time lag of Δ*t* = 3.4 s. We estimate the extrapolated solution corresponding to a vanishing grid size (Δ*x* → 0), where *e*12 and *e*23 represent the relative errors between meshes 1–2 and 2–3, respectively, and *e*^*extr*^ represents the relative error between the extrapolated solution and mesh 3. The close agreement between meshes 2 and 3 (*e*_23_ *<* 3% and *e*^*extr*^ *<* 1%) confirms mesh independence. Accordingly, we use mesh 3 for all subsequent simulations.

**Fig. 6.**
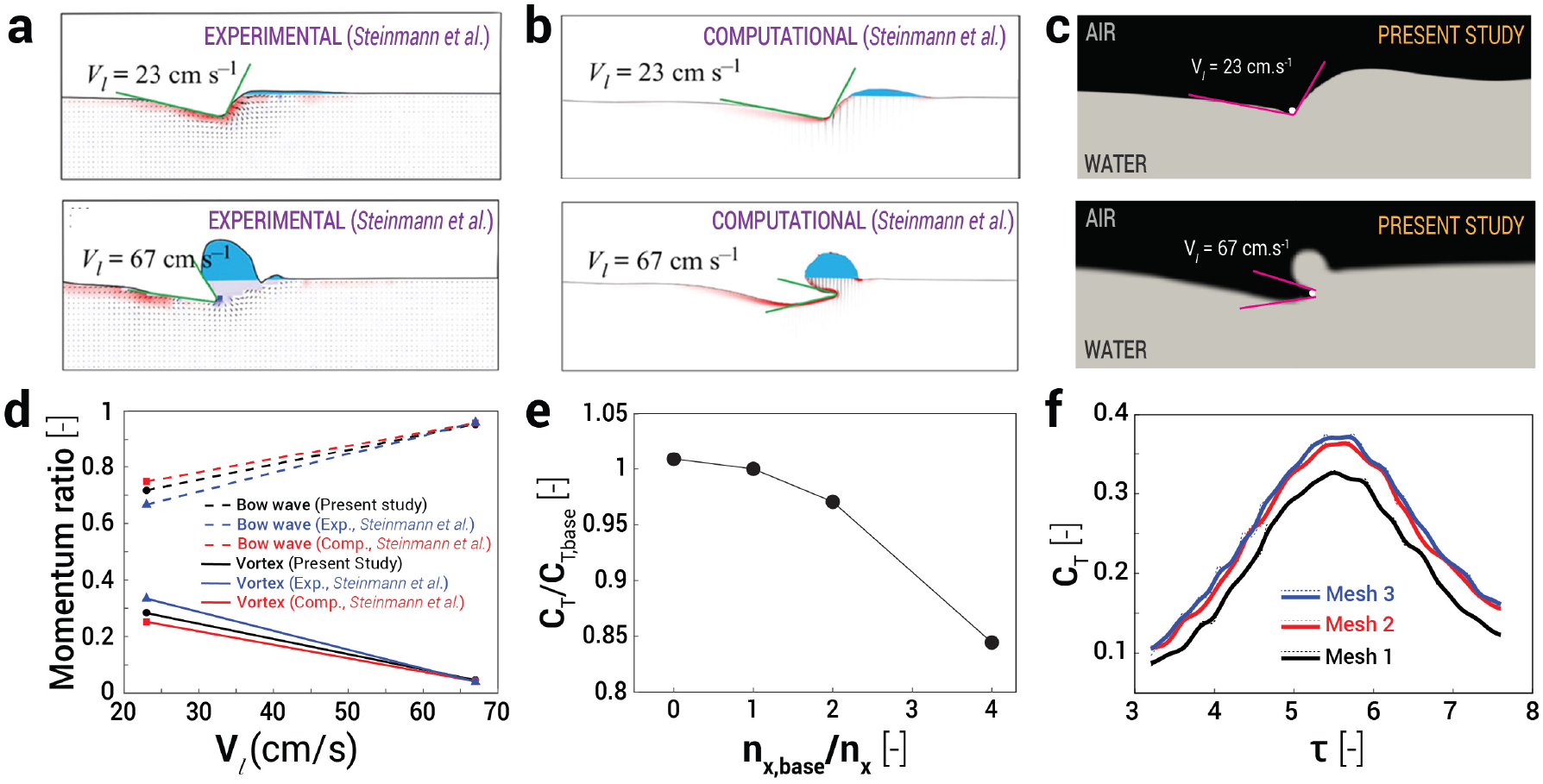
Validation and Verification of 2D two-phase CFD simulations. (a)-(d) Comparison between results from experiments and numerical simulations of Steinmann et al. (45) and the numerical model used for the present study. The sculling depth is kept fixed at *h*_*l*_ = 0.95 mm, and the leg velocity is kept at *V*_*l*_ = 23 cm/s for the first row and *V*_*l*_ = 67 cm/s for the second row. The quantities used for comparison are the bow wave momentum *p*_*bw*_, the time integral of surface tension forces *p*_*s*_, the anterior *α*_*a*_ and posterior *α*_*p*_ relative angles of the meniscus. (a) (First row) Experimental results (Steinmann et al., (45)) for *V*_*l*_ = 23 cm/s; *p*_*bw*_ = 2.8 × 10^*−*4^, *α*_*a*_ = 11 deg, *α*_*p*_ = 50 deg, *p*_*s*_ = 7.3 × 10^*−*4^. (b) (First row) Numerical results (Steinmann et al., (45)) for *V*_*l*_ = 23 cm/s; *p*_*bw*_ = 6.9 × 10^*−*4^, *α*_*a*_ = 11 deg, *α*_*p*_ = 50 deg, *p*_*s*_ = 7.3 × 10^*−*4^. (c) (First row) Numerical results (Present study) for *V*_*l*_ = 23 cm/s; *p*_*bw*_ = 1.9 × 10^*−*4^, *α*_*a*_ = 12.1 deg, *α*_*p*_ = 51.1 deg, *p*_*s*_ = 7.5 × 10^*−*4^. (a) (Second row) Experimental results (Steinmann et al., (45)) for *V*_*l*_ = 67 cm/s; *p*_*bw*_ = 2.3 × 10^*−*3^, *α*_*a*_ = 0 deg, *α*_*p*_ = 120 deg, *p*_*s*_ = 3 × 10^*−*3^. (b) (Second row) Numerical results (Steinmann et al., (45)) for *V*_*l*_ = 67 cm/s; *p*_*bw*_ =3.6 × 10^*−*3^, *α*_*a*_ = −14 deg, *α*_*p*_ = 166 deg, *p*_*s*_ = 4.2 × 10^*−*3^. (c) (Second row) Numerical results (Present study) for *V*_*l*_ = 67 cm/s; *p*_*bw*_ = 1.7 × 10^*−*3^, *α*_*a*_ = −13.9 deg, *α*_*p*_ = 160 deg, *p*_*s*_ = 4.1 × 10^*−*3^. (d) Comparison of the bow wave and vortex momentum ratios at different leg velocities derived from Experimental and Numerical results (Steinmann et al., (45)) and the numerical model used for the present study. (e) Results of grid sensitivity study conducted using Richardson extrapolation computed using 2D two-phase simulations for two cylinders undergoing prescribed 2D kinematics; *n*_*x*_ represents the cell count of the evaluated grids, *n*_*x*,*base*_ represents the cell count of Mesh 3, and *C*_*T*,*base*_ is the *C*_*T*_ computed from the Mesh 3. (f) Comparison of *C*_*T*_ variation with convective time *τ* (*τ* = (*tu*)*/d*) computed from 2D two-phase CFD simulations using three different cell count grids for two cylinders undergoing prescribed 2D kinematics.

We validate the present numerical model qualitatively by capturing the interface dynamics and quantitatively by comparing bow wave and vortex momentum ratios across various leg velocities, using the experimental and numerical results of Steinmann et al. (45) (Figure 6(a)–6(d)). Our numerical simulations are in close agreement with both the experimental and numerical results reported in their study.

We systematically studied the effect of submergence depth (*h*) and Reynolds number (*Re*) on the circulation of vortices shed from an isolated cylinder moving along a circular arc trajectory. In Figure 7, the evolution of vortices is shown with respect to the convective time *τ* (*τ* = (*tu*)*/d*). The vortex interactions remain qualitatively similar across the tested ranges of *h/d* = 0.25 to 0.75 and *Re* = 50 to 150 (Figure 7(a)-7(b) and Figure S13). As the cylinder grazes the air-water interface along its circular arc trajectory, a primary clockwise vortex gradually strengthens and reaches peak circulation just before the cylinder begins to exit the interface (see Figure S9 for circulation measurement details). The peak circulation of the primary vortex increased with both submergence depth and Reynolds number, as shown by the temporal evolution of circulation in Figures 7(c)–7(d). Side-view imaging of *Microvelia* walking on water informed our estimate of a shallow submergence depth of *h* = 0.5*d* for subsequent simulations. Note that this depth is much smaller than that observed in large water-walking organisms, such as Gerridae (45). For 2D multiphase simulations, we chose *Re* = 50 to balance computational cost with accurate resolution of the flow physics.

**Fig. 7.**
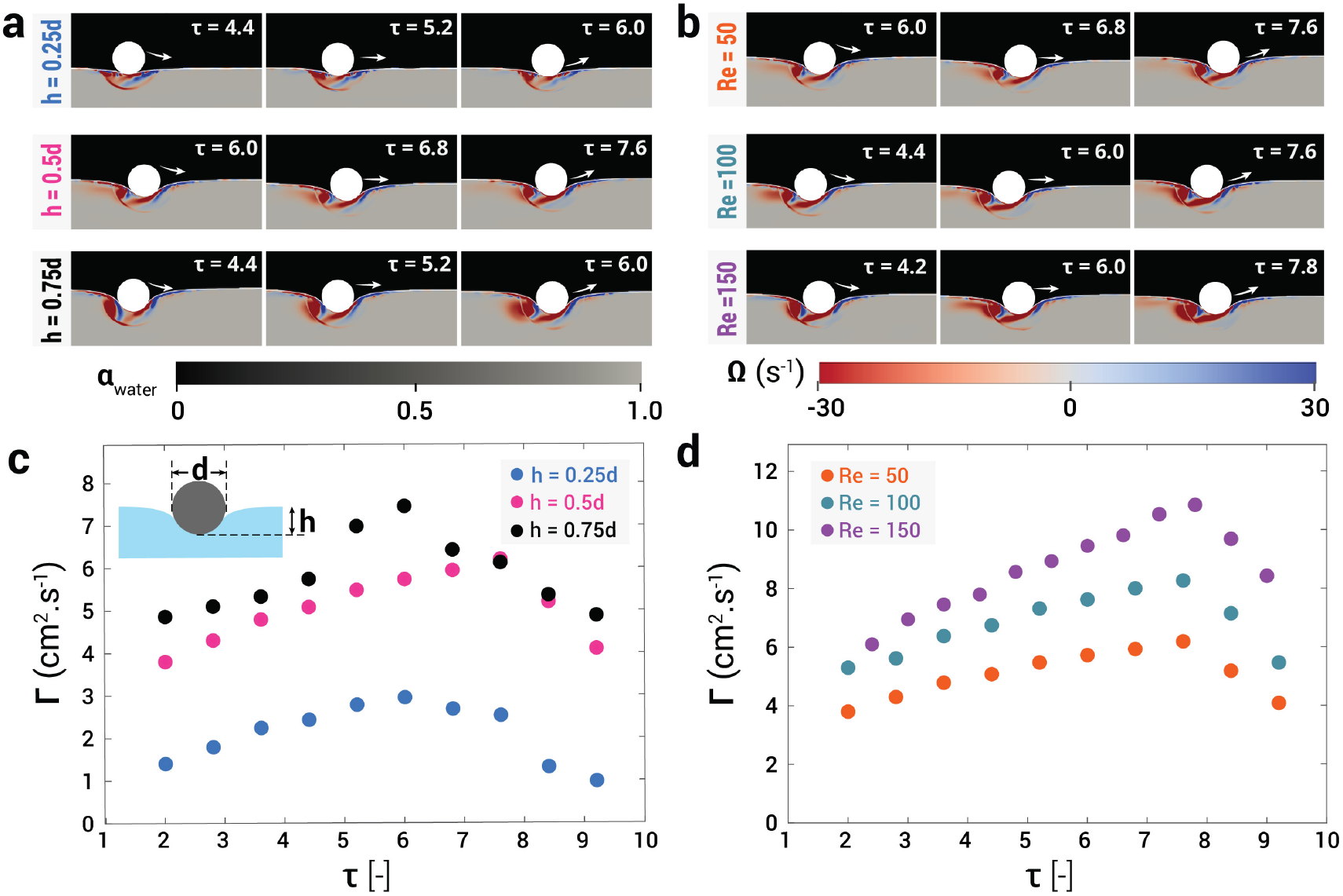
CFD results showing the effect of submergence depth and Reynolds number on the circulation of vortices shed from a cylinder moving along a circular arc trajectory. Temporal evolution of the primary vortex for different (a) *h* and (b) *Re* values; variation of Γ with convective time (*τ*) for different (c) *h* and (d) *Re* values. The different submergence depth cases are at Re = 50, and the different Re cases are at h = 0.5d. In the snapshots, the arrows indicate the direction of movement of the cylinder. As the cylinder enters the air-water interface, the primary vortex strength increases until it begins to come out of the water phase. The strength of the primary vortex shed from the cylinder increases with an increase in the submergence depth and the Re. This effect is qualitatively observed from the snapshots (a) and (b) and quantitatively from the circulation plots presented in (c) and (d).

To evaluate the effect of vortical interactions on thrust, we model the trajectories of the two cylinders on the water surface (Figure 8(a)) to mimic the side-view motion of *Microvelia*’s middle and hind legs with a time delay of Δ*t*. Figures 8(b)–8(d) show the temporal evolution of the primary vortex for three cases: single-body motion (no interaction), re-energization (Δ*t* = 3.4 s), and weak interaction (Δ*t* = 4.4 s). In the re-energization case, the clockwise interfacial vortex generated by the first cylinder constructively merges with a vortex with the same polarity from the second cylinder, enhancing both circulation and thrust. In contrast, in the weak interaction case, a longer Δ*t* causes the clockwise vortex from the first cylinder to interact with the counter-clockwise shear layer generated by the second cylinder, preventing constructive merging. As a result, the re-energization case shows significantly higher peak *C_T_* and corresponding peak Γ, while the weak interaction case shows lower values compared to the single-body case (Figures 8(e)-8(f)). The re-energization case shows an increase of ∼ 22.5% from *C_T_* =0.307 (single-cylinder) to *C_T_* = 0.376. Note that these 2D two-phase simulations are limited to only translation motion, and additional rotational motion at the interface, as observed for *Microvelia*, might result in a different magnitude of thrust enhancement.

**Fig. 8.**
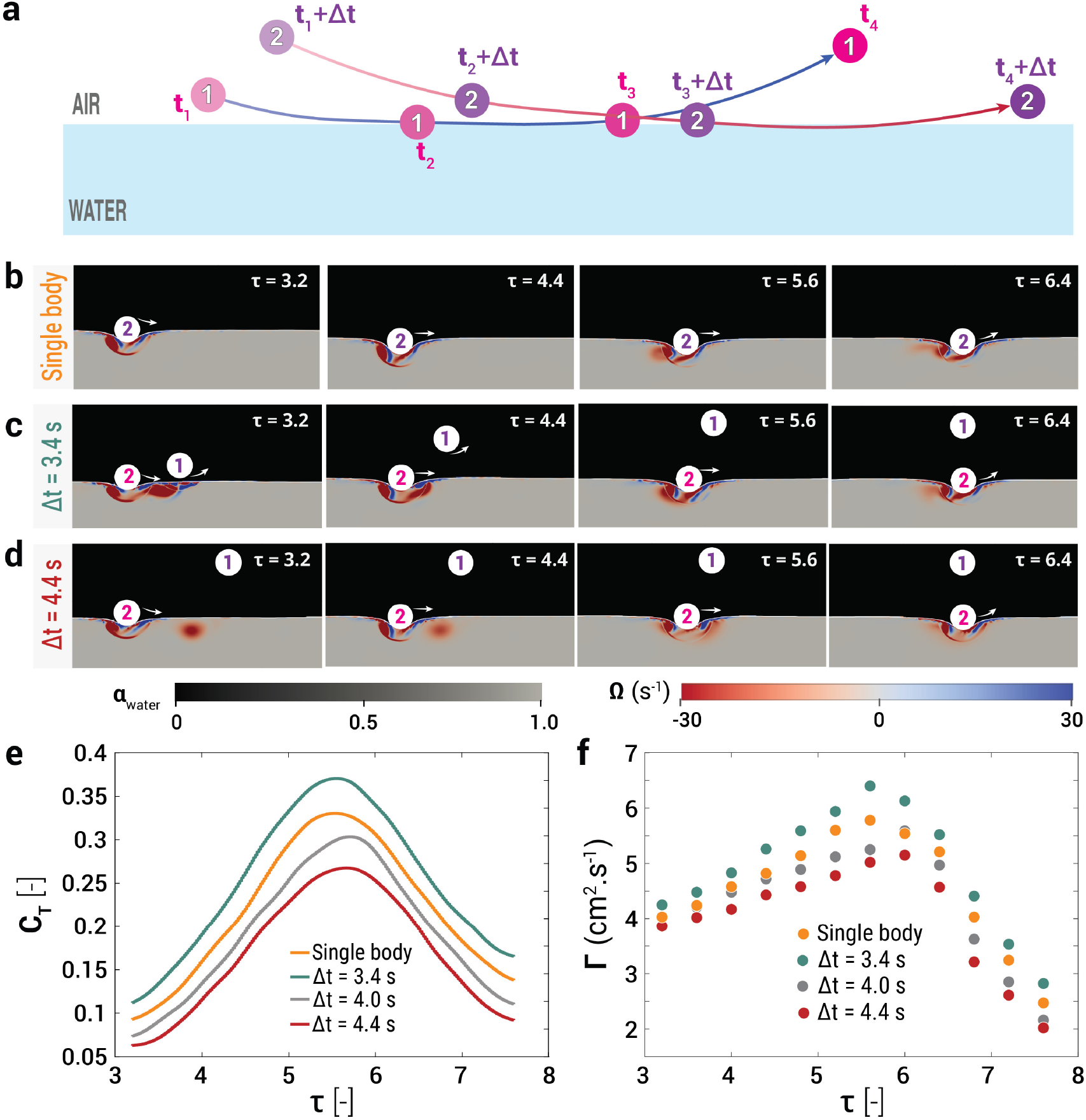
Comparison of the vortical interactions at the air-water interface and the resulting coefficient of thrust and circulation of the primary vortex for different time-delays between the motion of two cylinders for *Re* = 50. a) Schematic representation of the motion trajectories. First, cylinder 1 starts moving along its arc, and then, after a time delay of Δ*t*, cylinder 2 starts moving. This trajectory is representative of the middle and hind leg movement of the *Microvelia* tarsi. (b)-(d) Temporal evolution of the primary vortex shed from cylinder 2 moving along its trajectory. (b) No-interaction case with just cylinder 2 moving along its trajectory. (c) Re-energization case wherein cylinder 2 starts moving after a time delay Δ*t* = 3.4s and the vortex shed from cylinder 1 reinforces the primary vortex shed from cylinder 2. (d) Weak interaction wherein cylinder 2 starts moving after a time delay Δ*t* = 4.4s. The interfacial clockwise vortex generated by cylinder 1 ends up in the vicinity of the counter-clockwise shear layer generated on the second cylinder and prevents constructive merging with the same sense vortex. Temporal evolution of (e) *C*_*T*_, and (f) Γ with convective time (*τ*) of the second cylinder for the no interaction case, re-energization case (Δ*t* = 3.4s) and the weak interaction cases (Δ*t* = 4.0s and Δ*t* = 4.4s).

## Conclusions and Outlook

Our findings illuminate interfacial vortical interactions, the dynamic water-air boundary that supports diverse life forms. *Microvelia*, among one of the smallest and fastest water-walking animals, create nearly 2D vortices due to their minute size and weight, forming shallow dimples on the water surface (45, 47). Their alternating tripod gait, inherited from terrestrial ancestors, enables versatile movement across water, land, and duckweed (17, 32, 48).

Although less energy-efficient than the rowing gait, the alternating tripod gait excels in amphibious locomotion, providing *Microvelia* with a strategic advantage in foraging and evading predators (16, 24, 45). This gait and leg proportions facilitate interfacial vortex recapture combination, where the hind leg tarsi boost the circulation and fluid entrainment of vortices shed by the middle legs. This re-energization creates positive pressure at the hind tarsi, acting as a virtual wall that augments thrust (45). In other genera, such as *Mesovelia*, longer middle legs prevent effective vortex recapture, underscoring the critical role of leg size in this mechanism (Figure S10).

Interfacial vortex interactions hinge on the spatial location, angle of attack, and trajectory of hind leg tarsi, determining whether vortices are re-energized, weakened, or minimally interacted with. Our robotic arm physical model and CFD simulations reinforce the impact of inter-stroke intervals on these interactions. The data indicate that optimal timing and positioning of leg strokes enhance thrust through vortex re-energization, offering new insights into fluid dynamics at the air-water interface. However, thrust enhancement via vortex recapture is a localized phenomenon occurring at the tarsal scale; its implications for overall locomotion efficiency or energetic economy at the organism scale remain uncertain. Future studies involving juvenile *Microvelia* could clarify how body size and developmental stage influence vortex interactions. Extending our two-dimensional CFD analyses to three-dimensional multiphase simulations with empirical kinematics, and evaluating unsteady flow regimes representative of natural habitats, would further contextualize these findings. Synthetic robotic analogs recently developed by our team (49) could also provide a controlled experimental platform to directly assess how local interfacial vortex recapture affects locomotion performance at organism-relevant scales.

By uncovering the physics behind the vortical recapture in *Microvelia*, we extend similar mechanisms observed in jellyfish and fruit flies to the air-water interface (5, 11, 12). Interfacial vortex recapture could inspire the development of efficient water-skating devices and amphibious robots, enhancing our exploration of the oceanic and freshwater interfacial niches (26).

## Materials and Methods

### Sample Collection

*Microvelia Americana* (2-3 mm) were collected from freshwater streams and ponds located in Kennesaw and Austell in Georgia, USA (see Figure S1). They were kept in aquariums of dimensions 17.5in × 14in × 6.5in at a constant room temperature of ∼ 20^*°*^C with circadian lighting. *Microvelia* were fed live wingless fruit flies and/or springtails every day. Aquarium water was replaced every two weeks by removing about 20% of the water, as well as any debris, which was replenished with fresh water.

### Imaging and Image processing

*Microvelia* (*10* specimens) were filmed walking on water using a Photron FASTCAM Mini at 2000 frames per second (fps) with a resolution of 1024 × 1024. Nikon 70-200mm f/2.8G ED VR II AF-S Nikkor Zoom Lens was used to record the videos. *Microvelia* were placed on the water contained in square Petri dishes (*Thermo scientific*) of dimensions 10cm × 10cm × 1.5cm. The high-speed camera was vertically mounted to record the top view of the *Microvelia* locomotion on water. A LED light was placed under the Petri dish, separated by a light diffuser.

### Kinematics and statistical analysis

After recording the high-speed video, we used DeepLabCut pose estimation machine learning software to track several points along the body. The head and abdomen tips were tracked to digitize their time and position to calculate body speed and acceleration. The coxofemoral joint (hip) and the tarsal tip of each leg were tracked to calculate the stroke amplitude, and the coordinates of the tarsal tip were used to calculate leg speed, leg acceleration and stroke frequency. To calculate the Reynolds number (Re =**u***_t_d/*ν**), we used tarsal linear speed (**u**_t_), the diameter of the tarsus (d ∼ 50 *µm*), and the kinematic viscosity of water (**ν** = 1 ×10^−6^ m^2^/s).

Kinematic parameters, including stroke frequency, tarsal linear speed, and stroke amplitude, were measured and averaged for each individual run, consisting of 5 trials for 3 specimens each. Generalized Linear Mixed-Effects Models (GLMM) were fitted to the kinematics data, and pairwise comparisons were made to compare the statistical significance of each leg. To study statistical differences in relative impulse data from vortical interactions, we performed a Wilcoxon rank-sum test. Statistical analysis was performed using `glmer` and `wilcox` (R version 4.4.0) in the RStudio software (version 2024.04.1+748). Three statistical significance levels were used: ***(*p <* 0.001), **(*p <* 0.01), and *(*p <* 0.05).

### Particle Imaging Velocimetry

Particle Image Velocimetry (PIV) was used to visualize and measure the wake produced by *Microvelia* when walking on water. We seed the surface of the water with 1-5 *µ*m polystyrene particles (*ρ* = 1.3 g/cc, Copsheric, USA). A class-4 laser (Opto Engine LLC, 532 nm, 5 W) and a laser pointer (515-532 nm ± 5 Class IIIB, IVVTRYI, China) creates a laser sheet at the water interface to illuminate the particles for visualization and particle tracking (see Figure S3.a). *Microvelia* were then filmed moving along the surface at 2000 fps. The velocity and vorticity fields were estimated using PIVlab 2.61, a Matlab toolbox (43). Circulation of the vortices was measured frame-by-frame by selecting a constant area around the vortex pair in PIVlab.

### Pressure field reconstruction

The pressure field is estimated by numerical integration of the pressure gradient obtained from the PIV velocity field (figure S3.c). The surface flow is assumed to be locally two-dimensional, as the out-of-surface velocity is negligible. The first and second orders of velocity gradients are generated from the velocity field using the second-order finite difference method, from which the pressure gradient is calculated using the Navier-Stokes equation. To minimize the cumulative error of pressure gradient integration, the direction of integration is set to normal to the tarsi plane. The rectangular integration window is located at the tarsi center, and one side of this window is aligned with the tarsi plane. The window size contains 14 × 14 vectors sufficiently encompassing the vortex pair. To calculate the pressure difference, the data is extracted from two opposite lines parallel to the tarsi but 2 pixels away. The uncertainty of pressure is calculated from the error propagation theory as *δ*(*p*)*/p* ≈ 13.0%. The error from window selection is calculated as 18.0%.

### Physical model

We designed and built a physical model to understand the vortices’ interactions with the hind tarsi in *Microvelia*. The physical model consisted of two copper wires (*d* ∼ 0.1 mm) and two aluminum wires (*d* ∼ 1 mm), an Arduino Uno and two NEMA motors. The aluminum wires, powered by a NEMA motor, acted as two robotic arms. Thin copper wires attached at the front of the aluminum wires interacted with the water interface in a manner similar to that of *Microvelia* tarsi, shedding bipolar vortices in their wake. The stroke speed was controlled with an Arduino Uno. The duration, magnitude and interval between the two arms entering and leaving the surface of the water were regulated by altering the orientation of the aluminum wires.

### Computational Methodology

Two-dimensional laminar bulk phase and two-phase simulations of the low Reynolds number flow have been carried out by directly solving the incompressible Navier-Stokes equations using the finite volume method-based open-source library `OpenFOAM`. For conducting the simulations, a quiescent fluid condition is considered to match the experimental conditions present in the case of *Microvelia* and the physical model. An overset mesh framework (also known as the Chimera technique) is used, employing the `overPimpleDyMFoam` solver for bulk phase simulations and the `overInterDyMFoam` solver for two-phase simulations. The `overInterDyMFoam` solver implements the `Volume of Fluid` method to model the interface dynamics between two-phase immiscible fluids. In the overset framework, cell-to-cell mappings between the disconnected domain and body mesh regions are established, enabling complex body motions without the penalties associated with deforming meshes. The `inverse distance` overset interpolation technique is used to facilitate the interpolation between the `donor` and `acceptor` cells, while the cells inside the plates are considered to be holes, which do not take part in the calculations. The temporal and spatial discretization schemes are second-order accurate.

For the two-phase simulations, a wall contact angle (= 175^*°*^, (45)) is defined for the cylinders to model the superhydrophobicity effect present in the case of *Microvelia*. The kinematics of the cylinders are implemented using overset meshing. In the overset framework, cell-to-cell mappings between the disconnected domain and body mesh regions are established, enabling complex body motions without the penalties associated with deforming meshes. The `cellVolumeWeight` overset interpolation technique is used to facilitate the interpolation between the donor and acceptor cells, while the cells inside the plates are considered to be holes, which do not take part in the calculations.

For interface capturing, the `MULES` (Multidimensional Universal Limiter with Explicit Solution) algorithm is used, which is semi-implicit and second-order accurate in time, ensuring that the volume fraction of air and water remains bounded between 0 and 1. The Euler scheme is used for temporal discretization. The convective term in the momentum equation is discretized using a second-order, bounded linear Upwind scheme, while the convective term in the phase fraction transport equation is discretized using the Van Leer scheme, a second-order total variation diminishing scheme. Gradient discretization is carried out using the Gauss linear scheme with a cell-limited gradient limiter, and Laplacian terms are discretized using the Gauss linear scheme. Interpolation of cell center values to face centers is carried out using the linear scheme.

The present simulations utilize adaptive time stepping constrained by the maximum flow CFL number set to 1.0 and the maximum interface CFL number set to 0.5. For pressure velocity coupling, the `PIMPLE` algorithm is used, which is a hybrid between the `SIMPLE` (Semi-Implicit Method for Pressure-Linked Equations) and the `PISO` (Pressure-Implicit Split Operator) algorithms. In the present study, the `PIMPLE` algorithm uses two outer correction loops and three inner pressure correction loops. For the bulk phase simulations, a preconditioned conjugate gradient iterative solver is used to solve the pressure equation, whereas a diagonal incomplete-Cholesky method is used for preconditioning. A preconditioned smooth solver, with the symmetric Gauss-Seidel method as the preconditioner, is employed for solving the pressure-velocity coupling equation. The absolute error tolerance criteria for pressure and velocity are set to 10^−6^. On the other hand, for the two-phase simulations, the pressure equation is solved using the `PBiCGStab` solver with a DILU preconditioner, ensuring a tolerance of 10^−12^. The velocity is computed using the smoothSolver with a Gauss-Seidel smoother, with a residual tolerance of 10^−6^ and a maximum of 200 iterations. A `smoothSolver` with a Gauss-Seidel smoother is applied to the phase fraction terms, with a convergence tolerance of 10^−8^.

## ACKNOWLEDGMENTS

The authors thank the members of the Bhamla Lab for their feedback and useful discussions. SB acknowledges funding support from NIH MIRA Grant R35GM142588, NSF Grants CAREER 1941933 and PHY-2310691. PR acknowledges the funding support from the Eckert Postdoctoral Fellowship, Georgia Tech. JON acknowledges funding support from the GT UCEM fellowship program and the Herbert P. Haley fellowship program. CB acknowledges the support from EPSRC for the ARCHER2 Pioneer Project that provided the required computational resources. This work used the ARCHER2 UK National Supercomputing Service (https://www.archer2.ac.uk) for carrying out the simulations.

