## Supplementary material for "Interfacial vortex recapture enhances thrust in tiny water skaters": SI Info

#### This PDF file includes:

Figs. S1 to S13
Tables S1 to S3
Legends for Movies S1 to S4
SI References

#### Other supporting materials for this manuscript include the following:

Movies S1 to S4

### *Microvelia* habitat

Figure S1.a shows one of the examples of the habitat of *Microvelia americana*. These bugs are often found in low-flowing creeks, lakes, and ponds (1, 2). Insects such as ants and flies fall in the water and often struggle to swim out, becoming prey to water striders. Figure S1.c shows *Microvelia* trying to pierce its proboscis into an ant to feed on. *Microvelia* are often found to spend significant time on wet rocks within creeks; figure S1.d shows one of such instances.

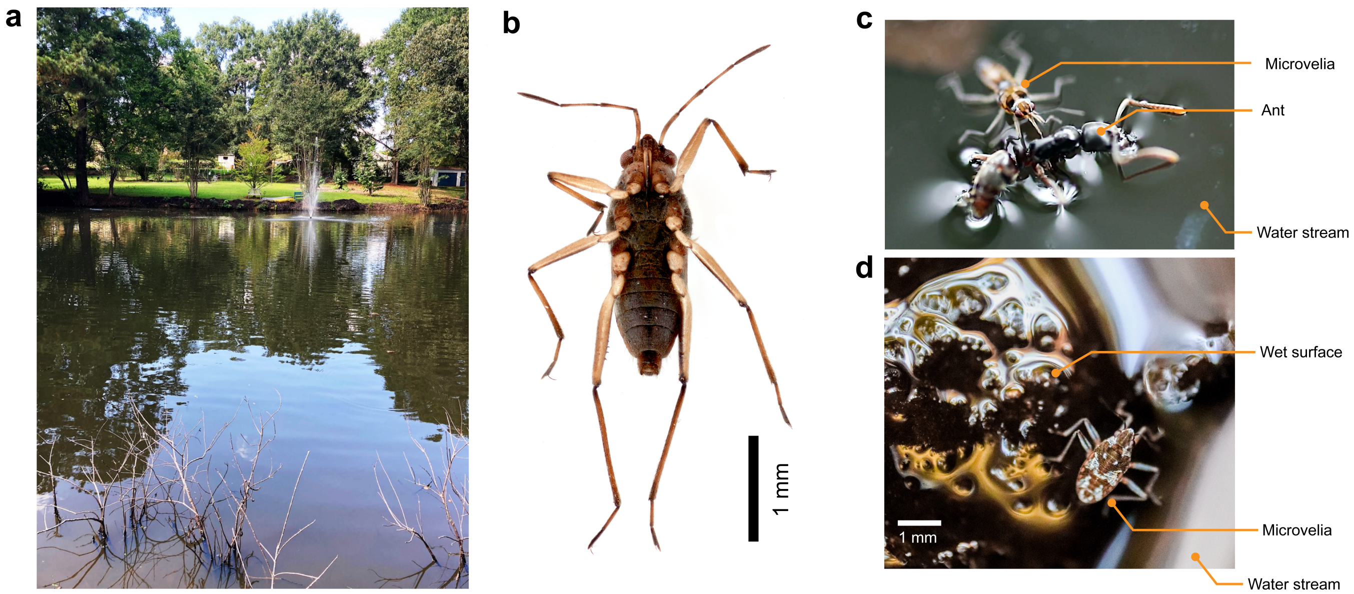


**Fig. S1. *Microvelia* in nature. (a).** A pond located in Austell, Georgia, USA, one of the source of *Microvelia americana* for our experiments. **(b).** Ventral view of *Microvelia americana*, **(c).** *Microvelia* using its proboscis to pierce the ant to feed on it, and **(d).** *Microvelia* on wet rock in a low-flow stream.

Figure S2 shows the variability in the entry points of the hind tarsi on the water surface relative to the exit points of the middle tarsi following their power strokes. This spatial variability contributes to differences in vortex circulation, the occurrence of weak interactions, and, in some cases, the hind tarsi completely missing the vortices shed by the middle tarsi.

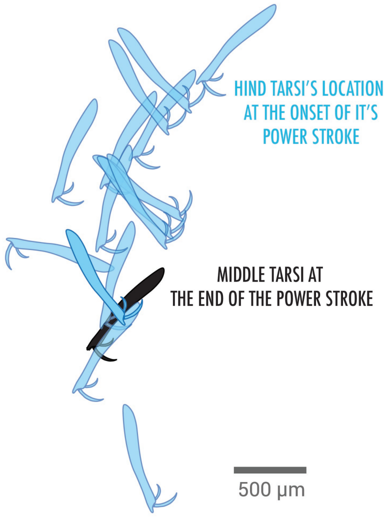


**Fig. S2. Positions of the hind tarsi at entry relative to the middle tarsi's exit point following the completion of its power stroke.**

### Particle Imaging Velocimetry

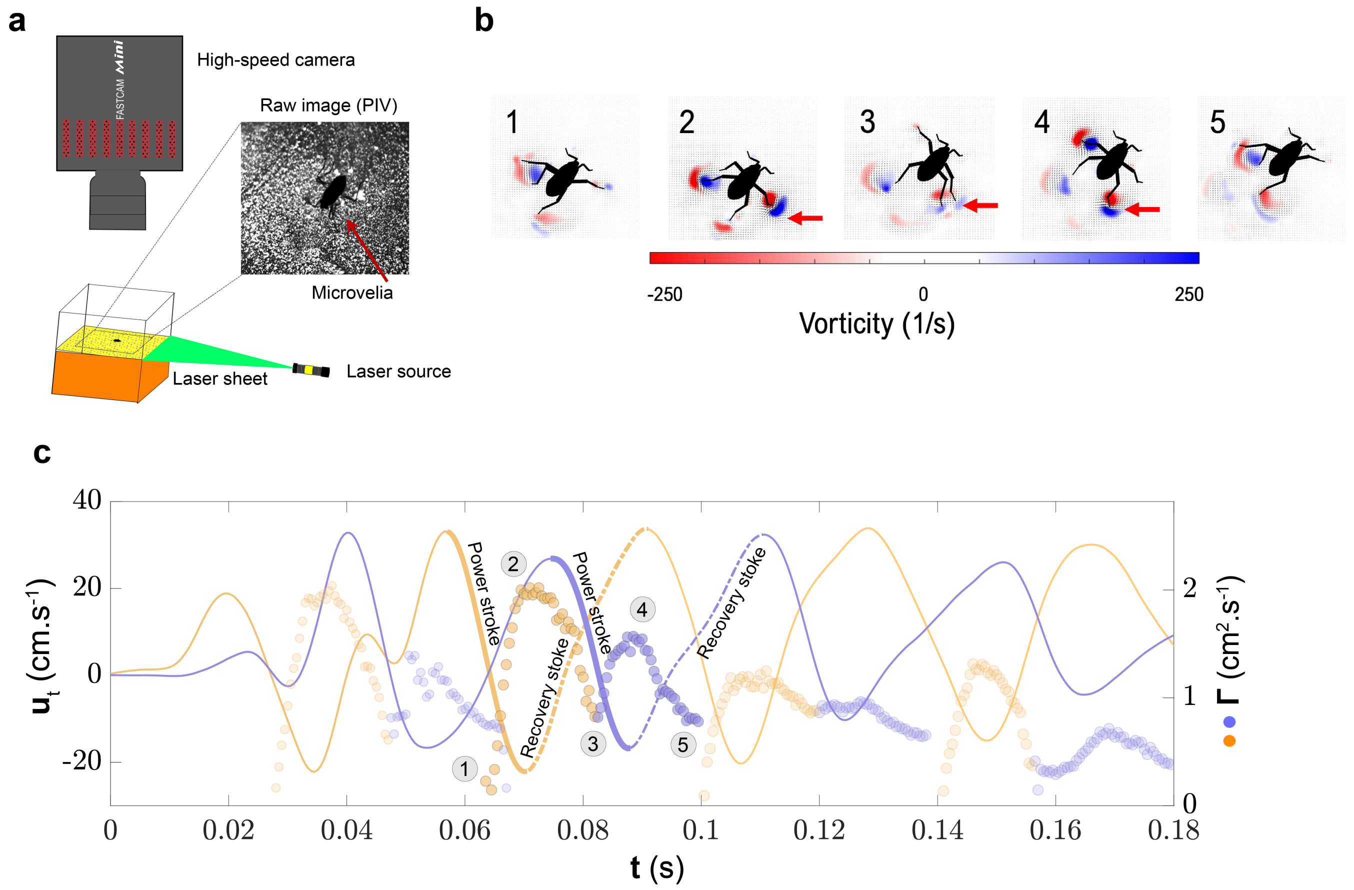


**Fig. S3. Particle Imaging Velocimetry (PIV) setup and results. (a).** Experimental setup to perform PIV to get flow field generated by *Microvelia* on water surface. **(b).** Correlation of tarsal speed and circulation of vortices generated by middle-right and hind-right tarsi stroking the water surface.

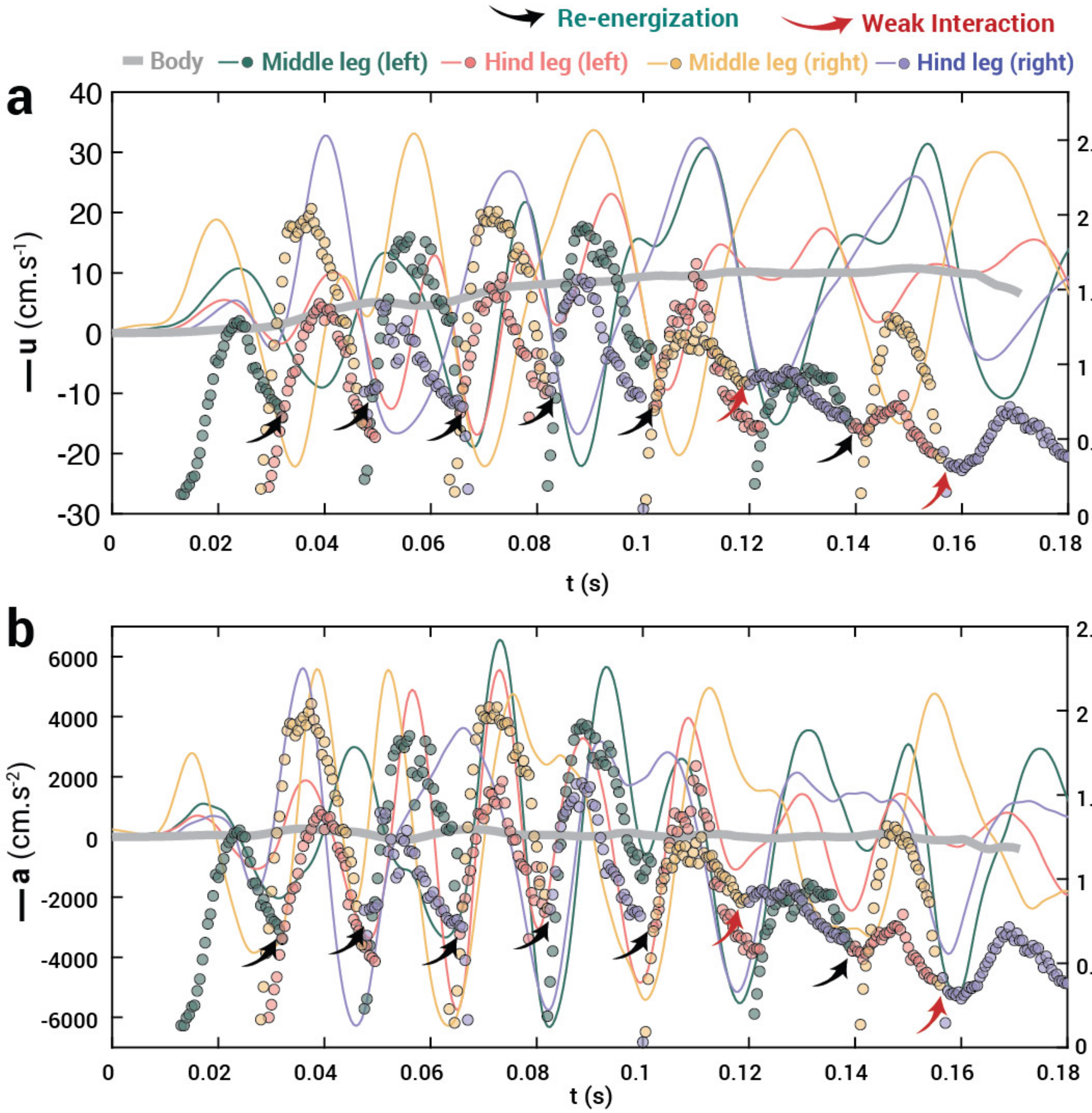


**Fig. S4.** Kinematics of *Microvelia* with circulation of vortices shed from its middle and hind legs. Temporal evolution of circulation with **a)** body speed and **b)** body acceleration.

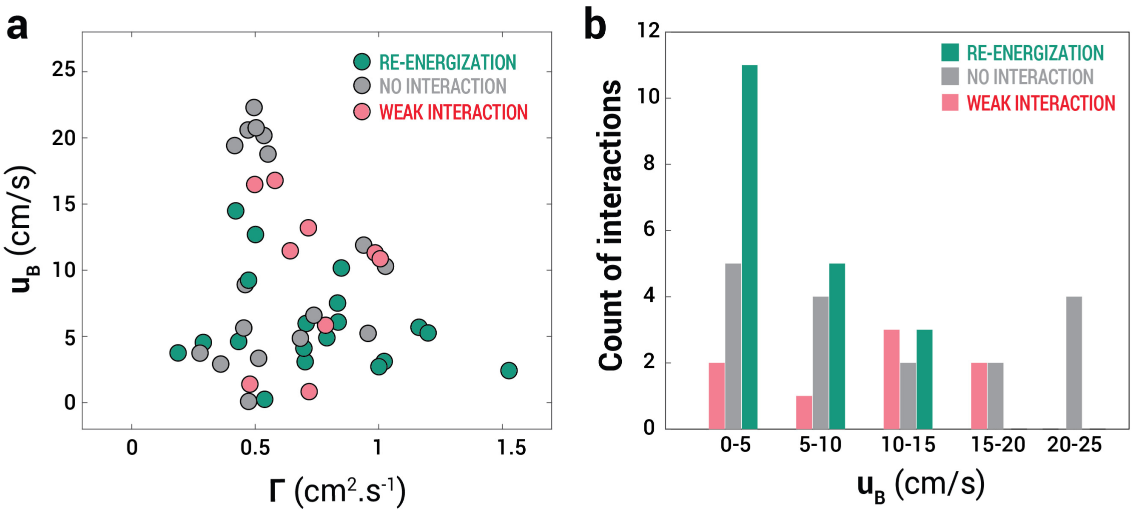


**Fig. S5. Dependence of vortical interactions on body speed. a)** Effect of *Microvelia*'s body speed on vortex circulation following interaction with the hind tarsi, showing that vortical re-energization predominantly occurs at lower body speeds. **b)** Histograms showing the frequency of each type of vortical interaction, further highlighting that re-energization is more common at lower body speeds. In contrast, no interaction is observed across a wide range of speeds, and weaker interactions tend to occur at body speeds up to 20 cm/s.

### Pressure Field Reconstruction

Pressure field was reconstructed using a custom Matlab script, where we estimate the relative impulse as an indicator of thrust enhancement resulting from vortical interactions. Pressure fields for different vortical interactions are presented in figure S6.

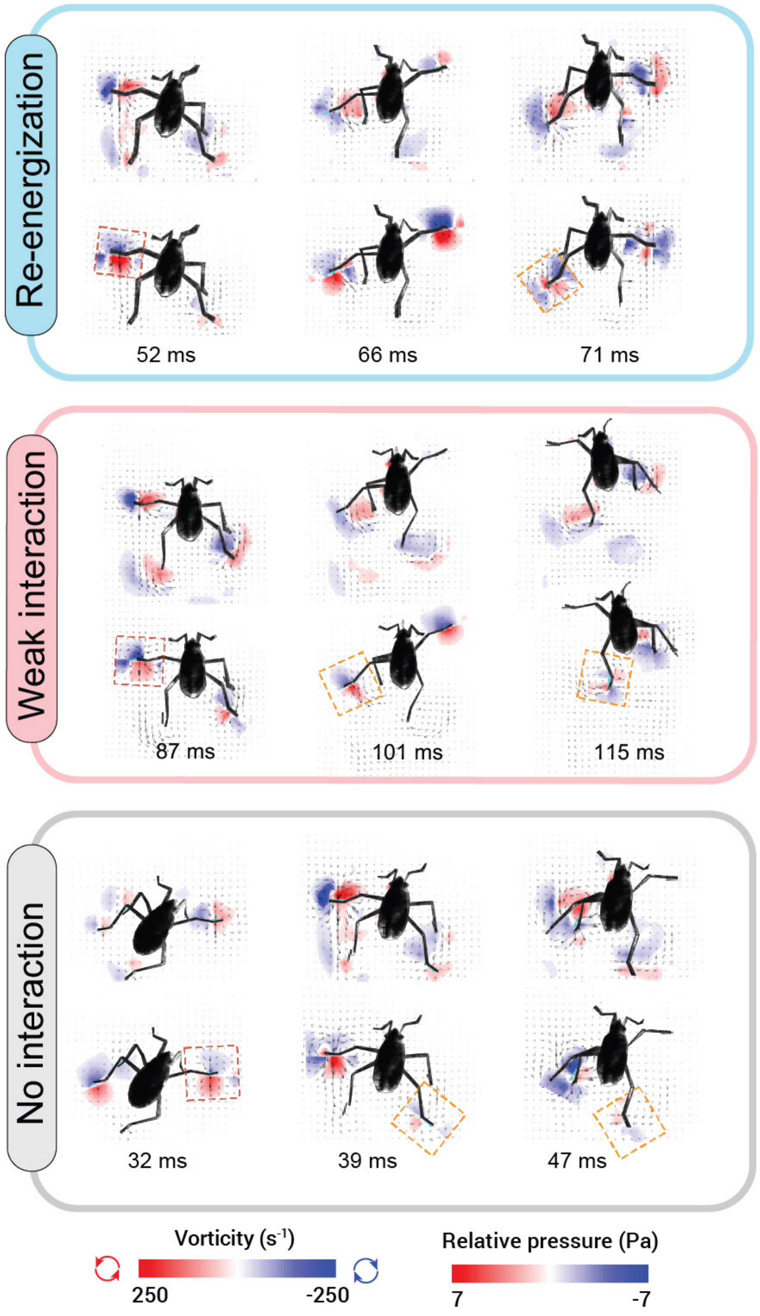


**Fig. S6. Pressure field reconstruction from the flow field data (PIV).** Representation of three cases of vortical interactions including (i) re-energization, (ii) no interaction, (iii) weak interaction. Top and bottom panels show vorticity and pressure fields, respectively for each case.

### Physical model

The physical model consisted of two robotic arms controlled by an Arduino Uno. The copper wires connected with robotic arms were made to skim the water surface, tracing the trajectories presented in figure S8.c. The outcome of vortical interactions depends on the inter-stroke interval of the two arms, $\Delta t_{21}$, which is the time interval between the onset of the stroke of the second arm and the completion of the stroke of the first arm. As the inter-stroke interval decreases, the second arm starts

 P. Rohilla[*,a], J. N. O'Neil[*,a], P. Singh[b], V. M. Ortega-Jimenez[c], D. Choi[a], C. Bose[d1], S. Bhamla[a]

interacting with the vortices from the first arm. Initially, the vortices were re-energized by the second arm until $\Delta t_{21}$ approaches zero, where vortices from both the first and the second arm interact weakly due to capillary waves generated by the entrance and exit of the copper wires to the surface of the water.

Increasing motor speed resulted in higher linear speed of the arm on the water surface, which created vortices with higher circulation (figure S8(a-d)).

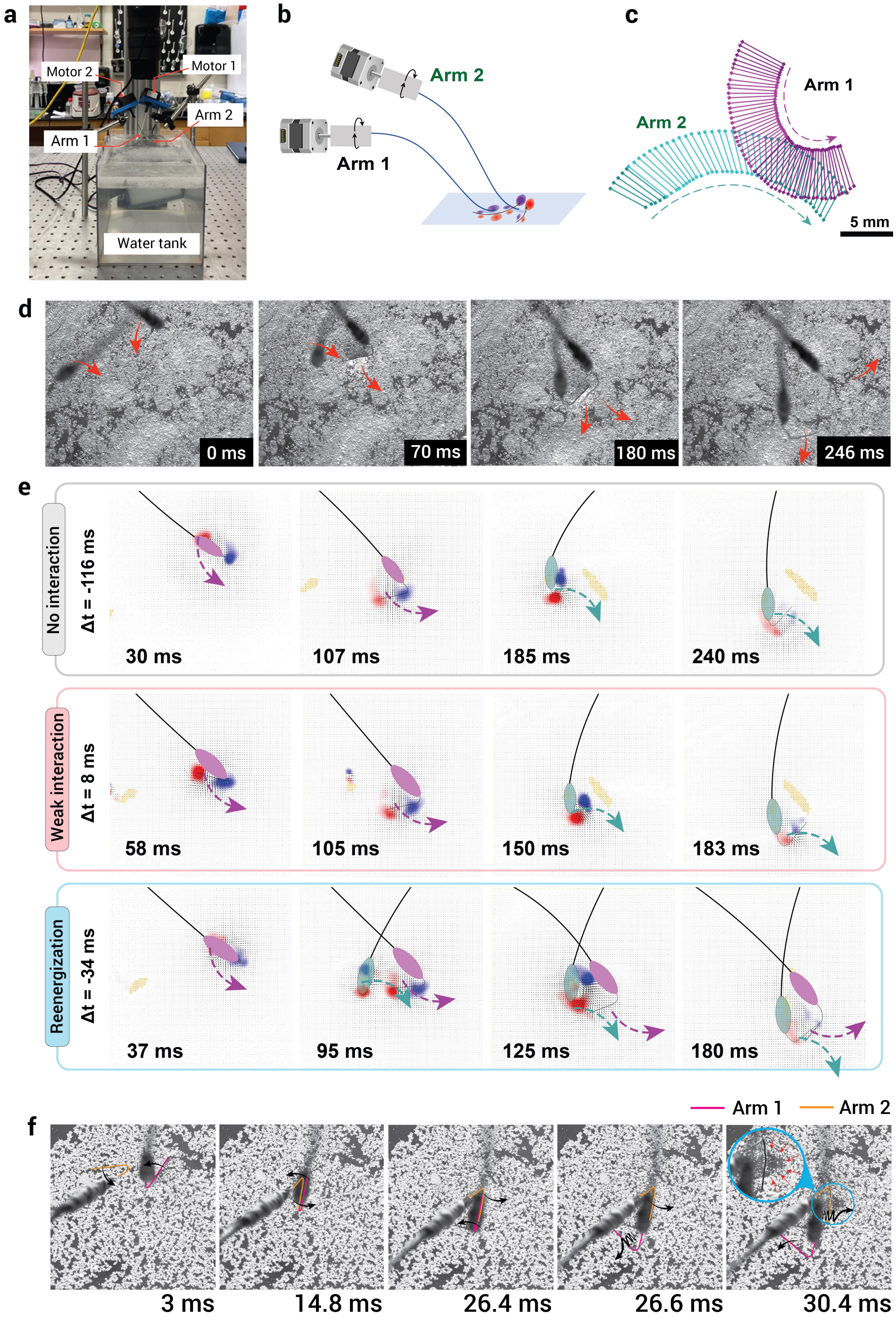


**Fig. S7. Physical model. (a).** Picture of the physical model installed over a water tank with high speed camera mounted vertically downwards. **(b).** Robotic arms stroking at the water interface. **(c).** Trajectory of two robotic arms while on water. **(d).** Snapshots showing robotic arms stroking the water surface. **(e).** PIV snapshots showing robotic arms shedding vortices and interacting with the vortices for varying inter-stroke intervals. **(f).** Collision of robotic arms beyond $\Delta t >$ 10 ms.

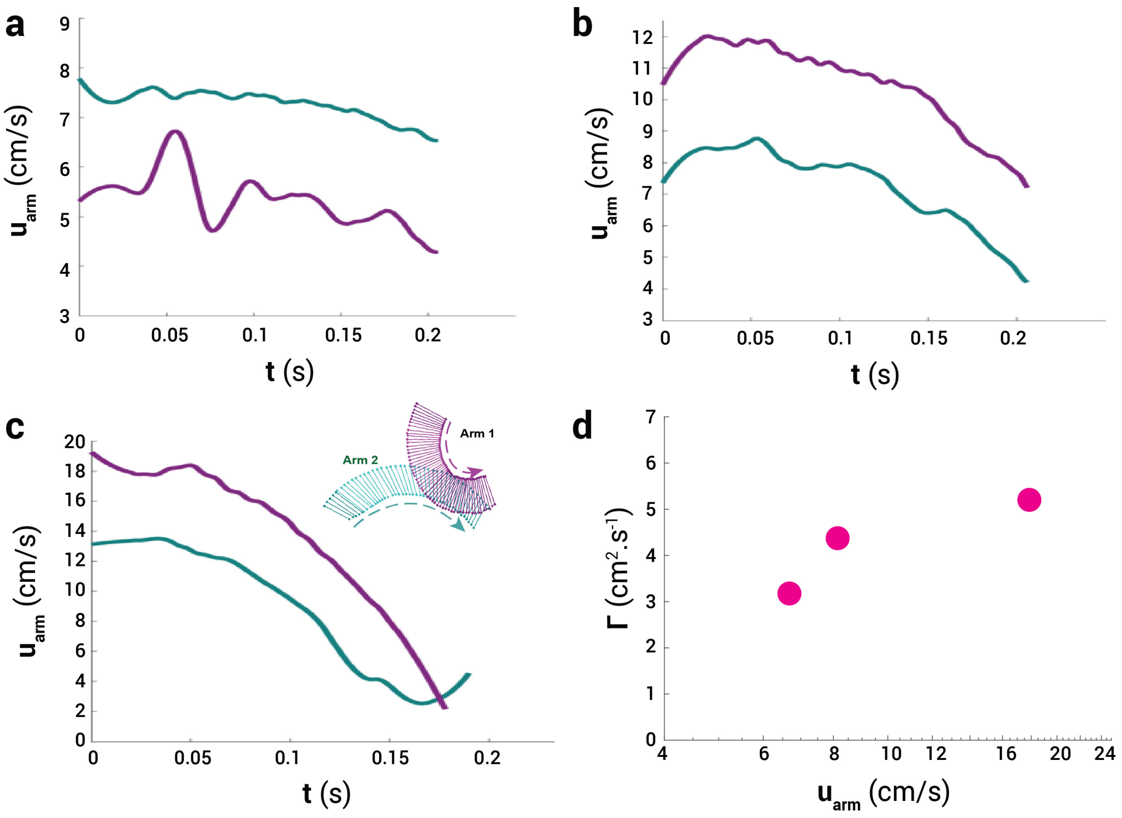


**Fig. S8. Arm's linear speed on water during skating for three different motor speeds (a-c). (d).** Effect of arm speed on the circulation of shed vortex pairs.

### Circulation measurement

We measured circulation in PIVlab (MATLAB) using a series of circular paths to compute the loop integral of tangential velocity around each selected circle, capturing the circulation of the enclosed vortex (see overlaid circles on the blue vortex in figure S9). As the radius of the circles increases, the measured circulation reaches a plateau; we use this saturated value as the representative circulation of the vortex at each time point. Additionally, we calculate the average circulation by averaging the magnitudes of circulation from both vortices in a vortex pair.

Similarly, in the 2D multiphase simulations, we measured vortex circulation by placing a series of concentric circles on the vortex behind the cylinder and selecting the maximum value as the representative circulation.

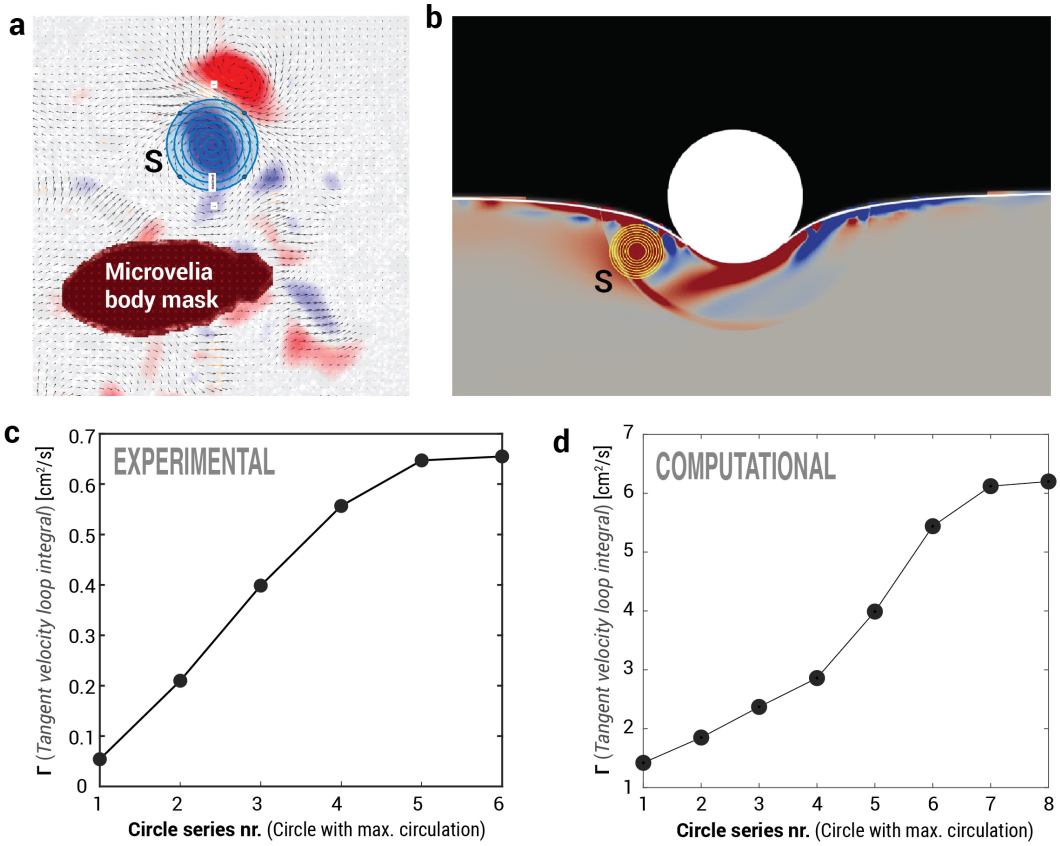


**Fig. S9. Circulation measurement.** Selection of circular regions to estimate the circulation within a vortex-bounded area for **(a)** *Microvelia* and **(b)** CFD. Circulation values for each circle, showing saturation at the maximum circulation of the vortex for **(c)** the 5th and 6th circles in the *Microvelia* case and **(d)** the 7th and 8th circles in the CFD case for a representative example.


P. Rohilla[*,a], J. N. O'Neil[*,a], P. Singh[b], V. M. Ortega-Jimenez[c], D. Choi[a], C. Bose[d1], S. Bhamla[a]

### Error analysis for reconstructing pressure field

We used Navier–Stokes equation for reconstructing pressure from PIV-based velocities:

$$-\nabla p \;=\; \rho\,\frac{\partial \mathbf{u}}{\partial t} \;+\; \rho\,(\mathbf{u}\cdot\nabla)\mathbf{u} \;-\; \mu\,\nabla^2\mathbf{u}.$$

The summary of error analysis for pressure field is as follows:

The velocity component $u_{\rm piv}$ is given by $u_{\rm piv} = M\,\Delta s/\Delta t$, where $M$ is the magnification factor, $\Delta s$ is the particle displacement, and $\Delta t$ is the time between image captures. The uncertainty is as follows (Lawson et al., 1999):

$$\frac{\delta(u_{\rm piv})}{u_{\rm piv}} \;=\; \sqrt{(\tfrac{\delta(M)}{M})^2+(\tfrac{\delta(\Delta s)}{\Delta s})^2+(\tfrac{\delta(\Delta t)}{\Delta t})^2}.$$

Here, the magnification rate (M) is $16.53\,\mu m$/px and maximum $\delta(M)$ resulted from reading error of scale bar is $0.22\mu m$/px. The error involved in the time interval, $\delta(\Delta t)$, is assumed as the inter-frame time interval was 1 $\mu s$ and $\Delta t = 1/2000s$. $\delta(\Delta s) \approx 0.04px$ given from the validation of cross-correlation algorithm (Scarano, 2001) and the max displacement of particles is $\Delta s \approx 4px$, the total velocity uncertainty is about $\delta(u_{\rm piv})/u_{\rm piv} = 1.7\%$. Since the maximum of $u_{\rm piv} \approx 0.3m/s$, $\delta(u_{\rm piv}) = 0.00495m/s$.

Approximating $\partial\mathbf{u}/\partial t$ by a central difference of order two leads to a truncation error $\mathcal{O}(\Delta t^2)$; the random (PIV) error propagates as $\delta\big(\partial u/\partial t\big) \sim \delta(u)/\Delta t$. The term $\rho\,(\mathbf{u}\cdot\nabla)\mathbf{u}$ involves first spatial derivatives and products of velocities. A second-order spatial scheme has truncation error $\mathcal{O}(\Delta x^2)$, while the random error scales like $\delta(u)/\Delta x$. For $\mu\,\nabla^2\mathbf{u}$, a second-order difference for the Laplacian has truncation error $\mathcal{O}(\Delta x^2)$. The random error for second derivatives is larger, given as $\delta\Big(\partial^2 u/\partial x^2\Big) \approx \sqrt{6}\,\delta(u)/\Delta x^2$. Thus the total error of pressure gradient is

$$\delta(\nabla p) \;\approx\; \sqrt{[\rho\,\delta(u)/\Delta t]^2 \;+\; [\rho\,\delta(u)/\Delta x]^2 \;+\; [\mu\,\sqrt{6}\,\delta(u)/\Delta x^2]^2}.$$

Once $\nabla p$ is computed, the pressure is recovered by $p(\mathbf{x}) - p(\mathbf{x}_0) = -\int_{\mathbf{x}_0}^{\mathbf{x}} \nabla p\,\cdot\,d\boldsymbol{\ell}$. A simple line integral with segment length $\Delta s$ has local truncation error $\mathcal{O}(\Delta s)$. Random errors in $\nabla p$ accumulate across $n$ segments. If $\Delta s = \Delta x = 0.15\,$mm and $n = 5$ to 6 (where $n$ denotes the maximum number of summation for the trapezoidal integration), the total random error can grow by a factor of order $\sqrt{n}$ if the errors are uncorrelated, which gives expression of $\delta(p) = \sqrt{n}\Delta x\delta(\nabla p)$. Thereby the total error of pressure, $\delta(p)$, is estimated as $11.67Pa$. Considering the maximum pressure in the flow is $\sim 100Pa$ (estimation from $p \sim \rho_f u_{tarsi}^2$ also gives $\sim 90Pa$), the uncertainty of pressure $\delta(p)/p \approx 13.0\%$. As the reviewer pointed out, the window size for pressure integration is critical and sensitive. A window that is too large leads to the accumulation of numerical errors, while a window that is too small can distort the boundary conditions. To quantify this, we tested three different window sizes of 10, 14, and 18 pixels, resulting in maximum pressure differences at the middle leg of 73.0, 89.0, and 97.8 $Pa$, respectively. This corresponds to a maximum error from window selection of 18.0%.

### Vortical shedding in Mesovelia

In Mesovelia, the hind legs are relatively longer than the middle legs. This length difference is one of the factors that prevent the hind tarsi from interacting with the vortices shed by the middle legs. Additionally, the timing of their strokes plays a crucial role; the hind legs enter the water well after the middle tarsi have completed their power stroke. Consequently, there is no interaction between the hind tarsi and the vortices from the middle tarsi.

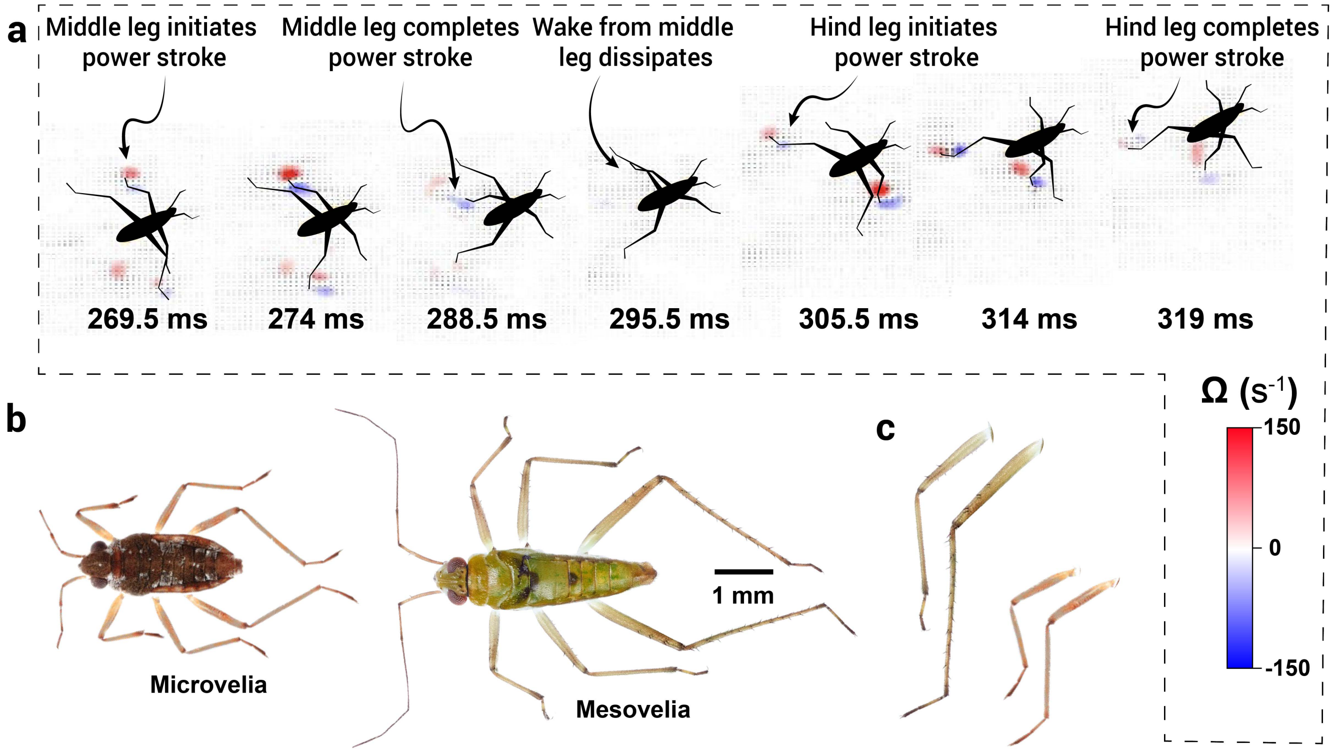


**Fig. S10. Vortical shedding in Mesovelia.** Snapshots showing the vortices shed by the left middle leg tarsus dissipates before the left hind leg enters the water. Moreover, the hind tarsi enters the water farther away being larger in length in comparison to the middle legs. Z-stack images of *Microvelia* and Mesovelia are presented side by side for their size comparison.

**Table S1. Dimensionless numbers for *Microvelia* locomoting on water**

| | Range |
|---|---|
| $Re$ | 2-21 |
| $Bo$ | 0.00021- 0.00057 |
| $We$ | 0.0014-0.0912 |
| $Ca$ | 0.0685-0.4384 |

 P. Rohilla*,a, J. N. O'Neil*,a, P. Singh[b], V. M. Ortega-Jimenez[c], D. Choi[a], C. Bose[d1], S. Bhamla[a]

### Vortical interactions in nature

From fruit flies capturing vortices shed by their wings (3) to jellyfish harnessing their own vortices for forward propulsion (4, 5), vortical interactions are ubiquitous in nature. Below in Figure S11, we present a comparison of these analogies of vortex capture in air and water to vortex capture at air-water interface in *Microvelia*.

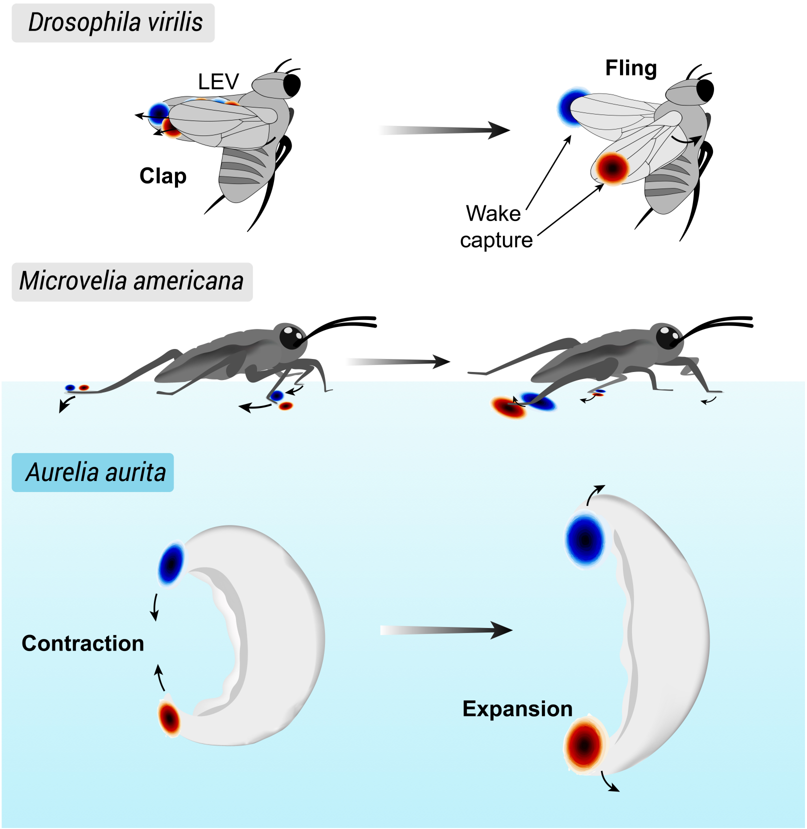


**Fig. S11. Comparison of vortical interactions across air, water, and the air-water interface.**

### CFD simulations

Mesh used in 2D single-phase and multiphase simulations are presented in figure S12. For 2D single-phase simulations, the first plate rotates and translates counterclockwise. The wake of plate 1 is then captured by the second plate, which rotates and translates clockwise, interacting with the wake of the first plate. These trajectories of the plates mimic the physical model arms' trajectory, as *Microvelia* tarsi trajectories could not be mimicked due to the spatial resolution of the biological data. The Reynolds number ($Re = 20$) is based on the plate thickness, which serves as an analog to both the cylinder diameter in the 2D multiphase simulation and the tarsi of *Microvelia*. For 2D multiphase simulations, cylinders follow an arc trajectory, entering - skimming the water surface and exiting the surface for Reynolds number ranging from 50-150.

Grid sensitivity test was conducted for 2D multiphase simulations and results corresponding to $\Delta t = 3.4$ s and Re = 50 are presented in table S2. Furthermore, we validated our simulation results with that presented in Steinmann et al. (6) (see table S3).

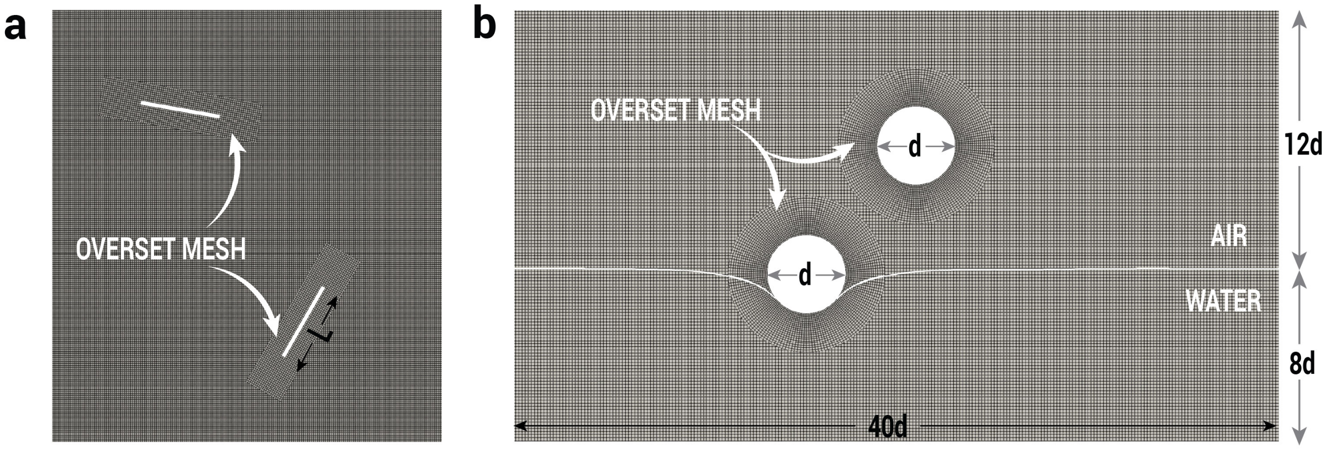


**Fig. S12. Meshes used in the CFD simulations. a)** Zoomed-in view of the CFD domain used for 2D single-phase simulations. A rectangular plate with length L and width L/20, representing a top view of a cylinder, is used to mimic the *Microvelia* tarsi. The original domain has a height and width of 30L. **b)** CFD domain used for 2D multiphase simulations. A circle with diameter d serves as an analog for the *Microvelia* tarsi. The domain has a length of 40d and a width of 20d. Note that the dimensions of these domains are not to the scale.

**Table S2. Results of grid sensitivity study for dt = 3.4s, Re = 50 case**

| Parameter | Value [-] | | | | $e_{12}$ [%] | $e_{23}$ [%] | $e^{extr}$ [%] |
|---|---|---|---|---|---|---|---|
| | Mesh 1 | Mesh 2 | Mesh 3 | Extrapolated Solution | | | |
| $C_T, Mean$ [-] | 0.2150 | 0.2471 | 0.2546 | 0.2569 | 12.99 | 2.94 | 0.89 |
| $\Gamma_{max}$ $(cm^2/s)$ | 5.79 | 6.28 | 6.40 | 6.44 | 7.80 | 1.88 | 0.60 |

**Table S3. Simulations parameters for *Gerridae* (6) and *Microvelia*.**

| | 2D 2-phase CFD (Gerridae) | 2D 2-phase CFD (Microvelia) |
|---|---|---|
| $d$ | 0.2 mm | 5 mm |
| $\mathbf{u}$ | 23 - 67 cm/s | 10 - 30 mm/s |
| $Re$ | 50 - 150 | 50 - 150 |

Snapshots of vortical interactions between cylinder 2 and vortex shed from cylinder 1 at Re = 150 is presented in figure S13 for $\Delta t$ ranging from 1.1-1.8 s. Similar trend in thrust coefficient ($C_T$) and circulation ($\Gamma$) was observed for 2D multiphase simulations corresponding to Re of 50 and 100.

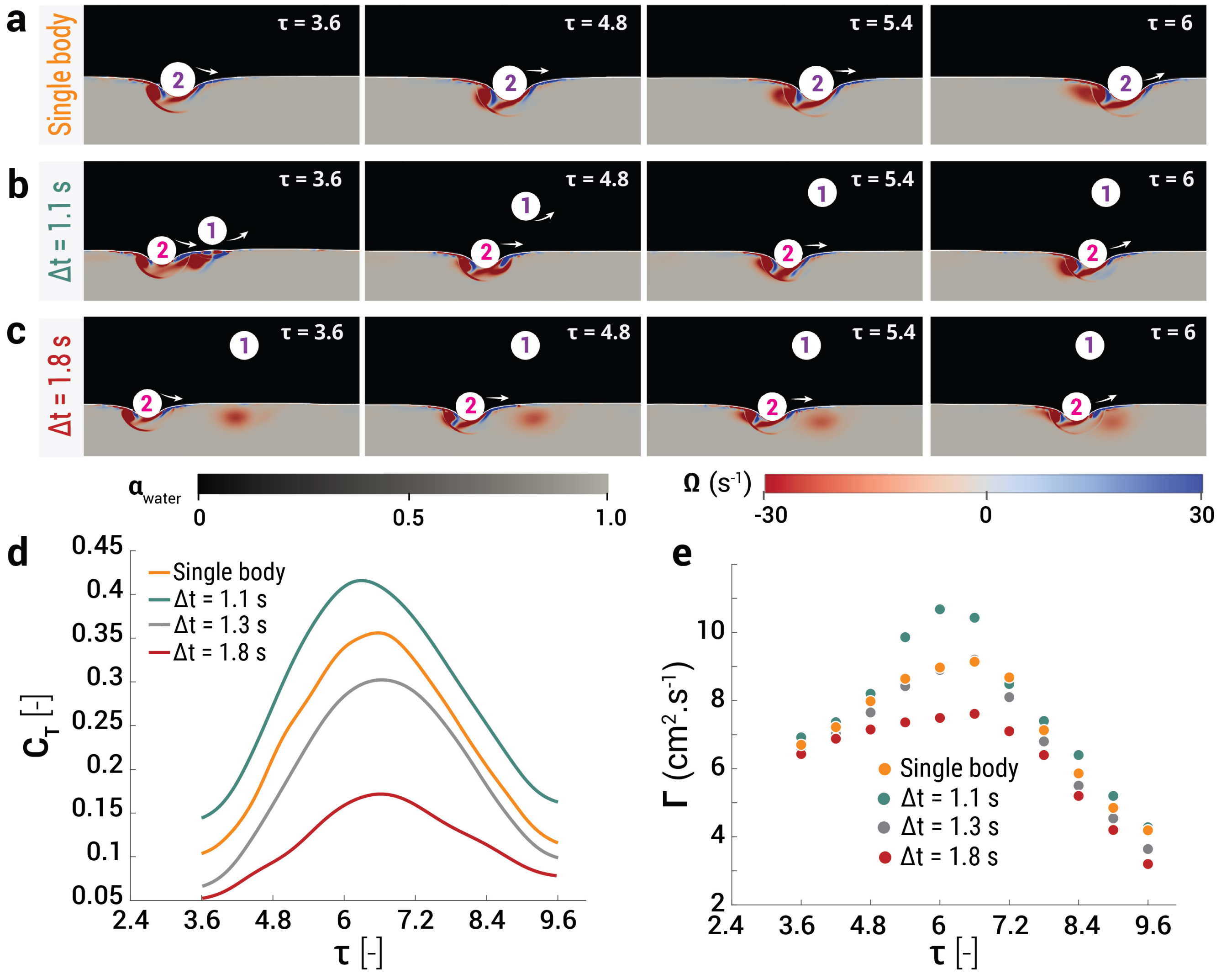


**Fig. S13. Comparison of the vortical interactions at the air-water interface and the resulting coefficient of thrust and circulation of the primary vortex for different time-delays between the motion of two cylinders for $Re = 150$.** a) No-interaction case with just cylinder 2 moving along its trajectory. b) Re-energization case wherein cylinder 2 starts moving after a time delay $\Delta t$ = 1.1 s and the vortex shed from cylinder 1 reinforces the primary vortex shed from cylinder 2. c) Weak interaction wherein cylinder 2 starts moving after a time delay $\Delta t$ = 1.8s. The interfacial clockwise vortex generated by cylinder 1 ends up in the vicinity of the counter-clockwise shear layer generated on the second cylinder and prevents constructive merging with the same sense vortex. Temporal evolution of (d) $C_T$, and (e) $\Gamma$ with convective time ($\tau$) of the second cylinder for the no interaction case, re-energization case ($\Delta t$ = 1.1 s) and the weak interaction cases ($\Delta t$ = 1.3 s and $\Delta t$ = 1.8 s).

**Movie S1. Biomechanics of *Microvelia*.**

**Movie S2. Vortical interactions in *Microvelia*.**

**Movie S3. Physical model reveals various kinds of vortical interactions.**

**Movie S4. Computational Fluid Dynamics modeling unveils the role of vortex capture on thrust.**
